# HuR-miRNA complex Activates RAS GTPase RalA to Facilitate Endosome Targeting and Extracellular Export of miRNAs

**DOI:** 10.1101/2023.06.22.546187

**Authors:** Syamantak Ghosh, Sourav Hom Choudhury, Kamalika Mukherjee, Suvendra N. Bhattacharyya

**Affiliations:** RNA Biology Research Laboratory, Molecular Genetics Division, CSIR-Indian Institute of Chemical Biology, Kolkata, India; Department of Pharmacology and Experimental Neuroscience, University of Nebraska Medical Center (UNMC), NE, USA

**Author notes:** Correspondence to be sent to or.

**Keywords:** miRNA, Endosomes, Endosome isolation, RNA import, RNA cargo trafficking, Extracellular vesicles, RalA GTPase, HuR, Rab5

## Abstract

Extracellular vesicles-mediated exchange of miRNA cargos between diverse types of mammalian cells is a major mechanism of controlling cellular miRNA levels and activity and thus to regulate expression of miRNA-target genes in both donor and recipient cells. Despite tremendous excitement related to extracellular vesicles-associated miRNAs as biomarkers or having therapeutic potential, the mechanism of selective packaging of miRNAs into endosomes and multivesicular bodies for subsequent extracellular export is a poorly studied area due to lack of assay system to study such processes *in vitro*. We have developed an *in vitro* assay with endosomes isolated from mammalian macrophage cells to follow miRNA packaging into endocytic organelles. The synthetic miRNAs, used in the assay, get imported inside the isolated endosomes during the *in vitro* reaction and become protected from RNase in a time and concentration dependent manner. The selective miRNA accumulation inside endosomes requires both ATP and GTP hydrolysis and the miRNA binding protein HuR. The HuR-miRNA complex binds and stimulates the endosomal RalA GTPase to facilitate the import of miRNAs into endosomes and their subsequent export as part of the extracellular vesicles. The endosomal targeting of miRNAs is also very much dependent on endosome maturation process that is controlled by Rab5 protein and ATP.

**Graphical Abstract:** 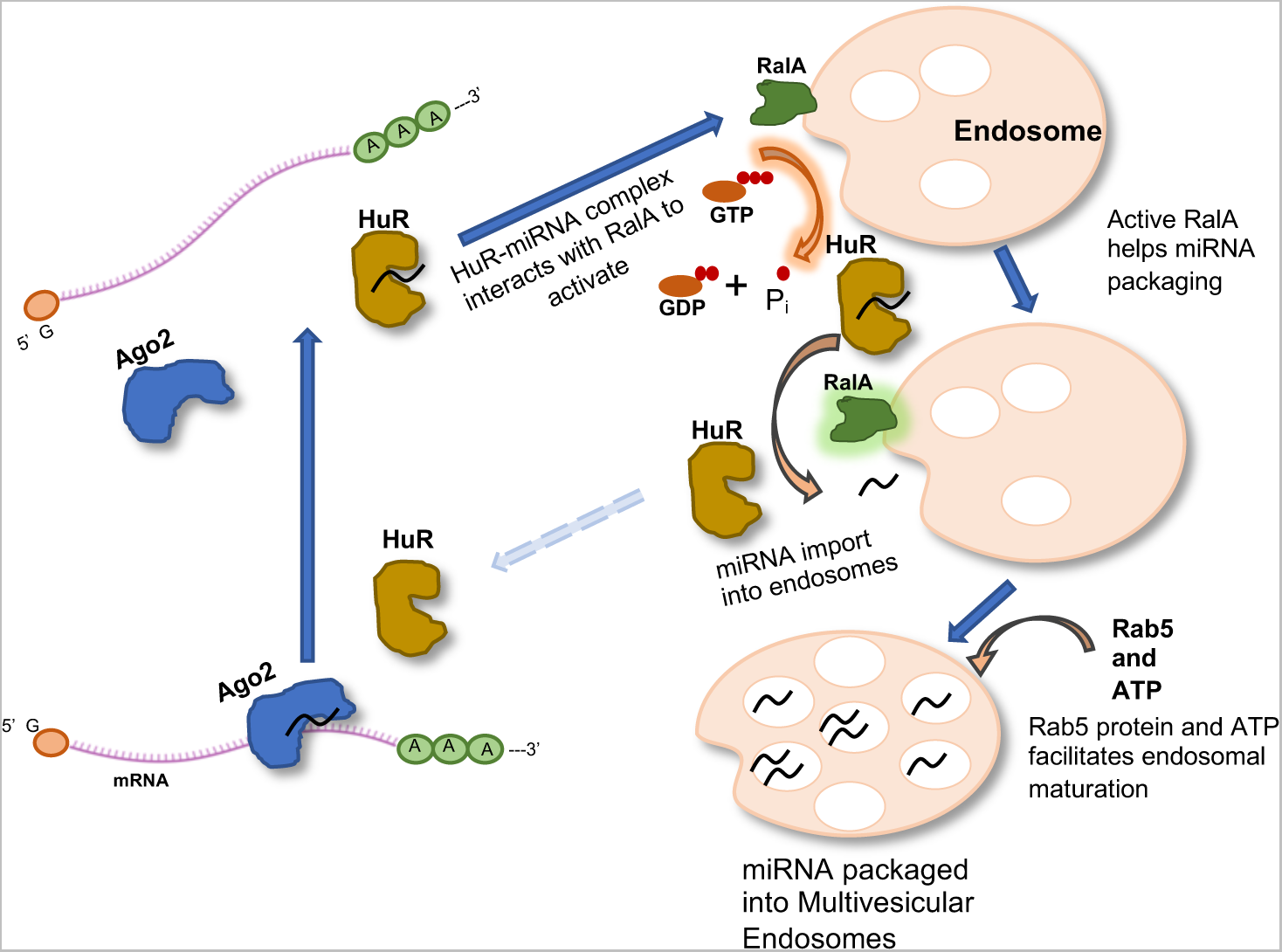

◦ miRNAs get imported to endosomes *in vitro and* are become protected to RNase
◦ Endosomal miRNA import is sequence specific and requires miRNA exporter protein HuR
◦ HuR-miRNA complex activates RalA GTPases to complete miRNA import.
◦ Rab5 protein and ATP hydrolysis is required for endosome maturation and miRNA import

## Introduction

MicroRNAs (miRNAs) are endogenous non-coding RNAs of 20-22 nucleotides length and are crucial modulators of post-transcriptional gene regulation in mammalian cells. miRNAs regulate gene expression either by repressing protein translation or by inducing degradation of miRNA-bound target mRNAs (Bartel, 2009). More than 2000 miRNAs are discovered in humans and about one third of human genes are regulated by miRNAs (Hammond, 2015). In body fluids, circulating miRNAs are found either in the free form or in association with Argonaute proteins or as part of extracellular vesicles (Arroyo et al., 2011; Kosaka et al., 2010).

Exosomes are one of the diverse types of extracellular vesicles (EVs) originated from endosomes or plasma membranes and released in the extracellular milieu. They are important players in intercellular communication by carrying cytosolic and membrane proteins, lipids and RNA within their lumen. Exosomes are formed in multivesicular bodies (MVBs), which upon maturation fuse with plasma membrane and release exosomes in extracellular space. The maturation of endosomes to MVBs requires cargo sorting at the endosomal membrane followed by membrane invagination and pinching of intraluminal vesicles (ILVs) inside the endosomal lumen to form the multivesicular structures (Raposo and Stoorvogel, 2013). There are evolutionary conserved multiprotein complexes known as- <u>e</u>ndosomal <u>s</u>orting <u>c</u>omplex responsible for transport (ESCRT)-0, -I, -II and -III which are associated with endosomes. ESCRT-0, -I and -II complexes recognize and sequester membrane proteins on the endosomal membrane, on the other hand ESCRT-III complex performs membrane budding and formation of ILV during endosome maturation (Hurley, 2010). However, ESCRT-independent mechanisms for MVB formation have been hypothesized in other reports (Buschow et al., 2010; Muralidharan-Chari et al., 2010; Nabhan et al., 2012; Stuffers et al., 2009; Trajkovic et al., 2008).

Sorting of miRNA to MVBs, for their subsequent release as part of exosomes, is an underexplored research area that has immense importance in EV-mediated intercellular communication and miRNA exchange. There are few published reports on importance of specific factors involved in this sorting process. RNA binding proteins such as, sumoylated hnRNPA2B1, SYNCRIP, Alyref and Fus have been identified to play a role in sorting of miRNAs into exosomes by recognizing specific short *EXOmotifs* present on exportable miRNAs (Garcia-Martin et al., 2022; Santangelo et al., 2016; Villarroya-Beltri et al., 2013). Additionally, in separate contexts, changes in dynamic transcriptome and KRAS overactivity may also regulate miRNA sorting into MVBs (McKenzie et al., 2016; Squadrito et al., 2014). RNA binding protein YBX1 was found to specifically sort miR-223 into exosomes by liquid-liquid phase separation (Liu et al., 2021; Shurtleff et al., 2016). In stressed human hepatic cells, HuR protein reversibly binds and accelerates export of miR-122 via exosomes to reduce the cellular miR-122 content to prevent cell death (Mukherjee et al., 2016). Role of HuR in sorting of other miRNAs such as miR-155 and let-7a has also been reported (Goswami et al., 2020). Recently, Lupus La protein found to be capable to export miR-122 because of its high affinity towards motifs present on miR-122 (Lang et al., 1988).

Even though involvement of these factors has been reported in miRNA sorting process, the exact mechanism and the machinery by which miRNAs, as cargo, get sorted into MVBs still remains elusive. In this study we aimed to identify internal factors or physiochemical parameters which drive miRNA into the lumens of multivesicular bodies. We have established an endosome enrichment protocol and have developed a cell-free assay by using endosome rich fractions from lysates of C6 glioblastoma cells to understand and validate several physiochemical factors which govern the miRNA packaging process in endosomes. Using this standardized cell-free assay we validated the efficacy of this system to import miRNA into endosomes. We observed ATP and GTP dependent endosome maturation and miRNA import which are controlled by miRNA-binding protein HuR and the small GTPase RalA. Our findings suggest interaction between HuR and RalA proteins that ensures HuR-miRNA complex-mediated activation of GTPase activity of RalA essential for miRNA packaging into MVBs and subsequent export via exosomes. The maturation of endosomes into MVB found to be also essential for endosomal uptake and export of miRNAs. Rab5, an early endosome associated protein, found to promote export of miRNA while its mutant defective for endosomal maturation failed to do so.

## Results

### *In vitro* import of single stranded miRNAs into endosomes isolated from mammalian cells

Association of miRNA-repressed mRNA and Ago2-miRNA complex have been detected with early endosome membrane (Janas et al., 2012).This association precedes miRNA dissociation from Ago2 and its packaging into endosomes for its export via EVs (Bose et al., 2020; Groot and Lee, 2020; Iavello et al., 2016). Therefore, isolated early endosomes can serve as a very good system to study the miRNA packing process *in vitro*. We had planned to use endosomes isolated from mammalian cells to develop the *in vitro* assay to study miRNA packaging. To that end, C6 or HEK293 cell lysates were separated on a 3-30% Optiprep^TM^ gradient to separate the endosomal compartments from rest of the organelles. The western blot analysis of the Optiprep^TM^ fractions for different marker proteins confirmed the presence of endosomal protein markers in the top lighter fractions of the gradients and that were separated from mitochondrial or lysosomal components (Fig 1A and B and Fig S1A). Subsequently, to separate the early endosomes from the late endosomes, we have used the 5-15% step gradient to pellet down the endoplasmic reticulum, mitochondria and lysosomes after centrifugation at 133,000xg for 2h. The separation of early from late endosome was evident in western blot analysis for early (Rab5, Alix) and the late endosomal protein (Rab7a). The pooled early endosomal fraction of the 5-15% gradient (Fractions 1-3) was western blotted for respective marker proteins. The early endosomal fractions were enriched for Alix, Rab5A but have reduced level of late endosomal protein Rab7a while ER marker protein Calnexin was absent in these fractions (Fig 1C and D and Fig S1B). We characterized the isolated organelles for their size and shape by AFM and NTA analysis to document the presence of organellar structures in the isolated fractions having the average size of 180-200nm and average concentration of 2.5X10^9^/ml (Fig1E and F). To follow the trafficking of miRNA into the endosomes and to understand the mechanistic detail of this trafficking, we used early endosome enriched fractions from HEK293 or C6 glioma cell lysates and used the isolated endosomes for the *in vitro* miRNA import assay (Fig 1G). In this assay, endosomes were incubated with phosphorylated form of single stranded miR-122 in the *in vitro* reaction and it was followed by an RNase treatment to degrade the residual or membrane attached miRNAs that were not getting trafficked inside the endosomes and thus were not protected from RNase after the miRNA import reaction. The miR-122 is not expressed in C6 or HEK293 cells and thus the isolated endosomes were free from pre-existing miR-122 pool. The RNase protected miR-122 pool was retrieved and quantified. The imported miR-122 level was normalized against the levels of miR-146a already present in the endosomes isolated from the C6 cells. Levels of miR-146a did not get altered during the assay (Fig S1F). We have observed a time dependent increase of miR-122 content in isolated endosomes till 30 mins of incubation with a drastic reduction in import level after 60 minutes of incubation possibly due to loss of vesicular integrity of the isolated endosomes during prolonged incubation of 60 mins (Fig 1H and Fig S1C). Incubation of endosomes with increasing concentration of RNase removed majority of residual RNA present in the reaction and only the imported RNA remained intact (Figure S1D). To confirm that only the membrane protected and imported RNA are RNase resistant after the import assay *in vitro*, endosomes were ruptured with deoxycholate before the RNase treatment but after the import reaction. The recovered membranes were depleted for miR-122 while Rab5 was found to remain associated with the ruptured membrane of endosomes (Fig 1J). Increasing concentration of substrate RNA also showed an increase in net number of imported miRNAs but the amount of imported miRNA reached saturation for a given amount of endosomes and does not increase significantly with further enhancement of substrate miRNA concentration (Fig1K and Fig S1E). miR-122 pre-loaded endosomes, isolated from C6 cells exogenously expressing miR-122 did not show any change in endosomal miR-122 content during the import assay. This result confirms the intactness of the reisolated endosomes after the import reaction of 30 mins and subsequent RNase treatment (Fig S1G). The import assays with other miRNAs as substrates were also done. With synthetic miR-146a and miR-155, we had documented selective import of miR-146a over miR-155 in the assays done *in vitro* (Fig 1L). Incubation of total small RNA pool isolated from C6 cells with endosomes followed by an RNase treatment and recovery of the protected RNA showed a differential import of different miRNAs present in the reaction mix and thus confirms a selective nature of the import machinery present on isolated endosomes (Fig 1M).

**Figure 1.**
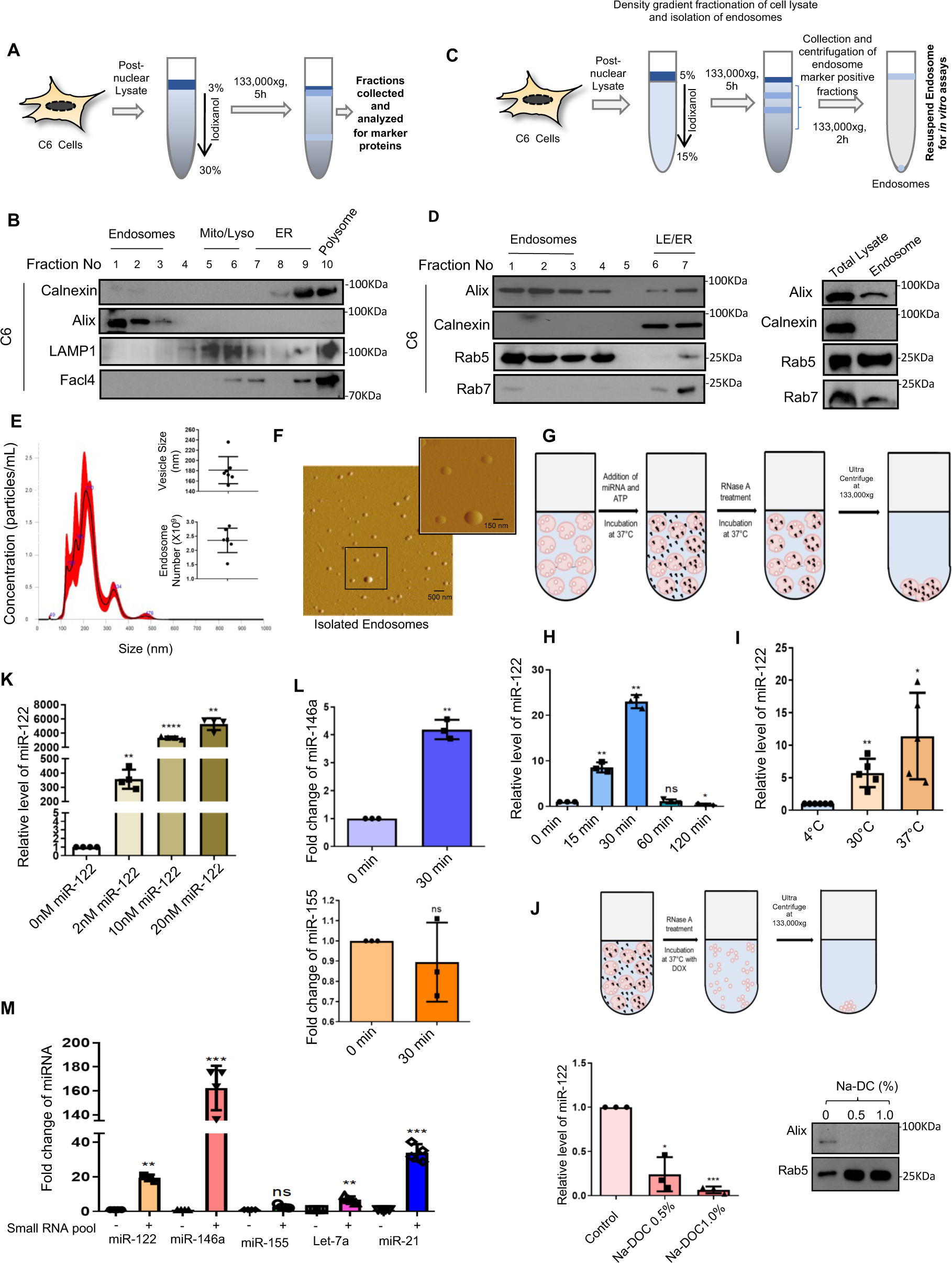
Endosomes isolated from mammalian cells import miRNAs in an *in vitro* assay. **A.** Schematic workflow of Subcellular fractionation of C6 cell lysate on 3-30% continuous iodixanol density gradient by ultracentrifugation and collection of fractions. **B.** Western blots of various organelle-specific marker proteins to detect in fractions obtained from subcellular fractionation of C6 post-nuclear lysate separated on 3-30% iodixanol gradient. **C.** Schematic workflow of subcellular fractionation of C6 cell lysate on 5-15% iodixanol step gradient by ultracentrifugation and collection of endosome rich fractions. **D.** Western blots of organelle specific marker proteins in different fractions obtained from 5-15% iodixanol step gradient fractionation of C6 post-nuclear lysate. The fractions 1-3 were pooled down to reisolate the endosomes as pellet and also analysed for marker proteins (right most lane). **E.** Size distribution of isolated endosomes in nanoparticle tracking analysis (NTA) and quantification of average size and concentration of endosomes analysed by NTA. **F.** Visualization of endosomes isolated from C6 cells by Atomic Force Microscopy (AFM)(Scale bar 500nm). A higher magnification image of endosomes is in the inset (Scale bar 150nm). **G.** Schematic representation of the cell-free reaction workflow to follow miRNA import into endosomes **H.** Relative amount of synthetic miR-122 (10nM) as cargo that get imported with increasing time of incubation in an import buffer with isolated endosomes followed by an RNase treatment to get rid of the non-imported miRNAs. The imported miRNA was quantified by qRT-PCR and value at 0 min of incubation was considered as unit. Endogenous miR-146 level present in endosomes were used for normalization (n=3 independent experiments, P= 0.007, 0.0014, 0.7143 and 0.043). **I.** Effect of temperature on miRNA import. The imported miR-122 was quantified by qRT-PCR and value at 4^0^C incubation temperature was considered as unit. Endogenous miR-146a level present in endosomes were used for normalization (n=5 independent experiments, P= 0.0084, 0.0249) **J.** Imported miRNAs are protected from RNase. After the import reaction increasing concentration of ionic detergent sodium deoxycholate was added to rupture the endosomes and expose the imported miR-122 to Rnase A (100ng/µL) for degradation (upper panel). The level of miRNA with or without detergent treatment of endosome were removed and analysed by qRT-PCR. Endogenous miR-146a present within endosomes were used for normalization (n=3 independent experiments, P= 0.021, 0.0006; lower left panel). Immunoblot of RNase and detergent treated vesicles showing amount of endosomal protein Rab5 that was not solubilized by the detergent from endosomal membrane while Alix was extracted with the detergent applied (lower right panel). **K.** Amount of miR-122 getting imported and protected from RNase with increasing amount of substrate concentration. The imported miRNA was quantified by qRT-PCR. Endogenous miR-146 level present in endosomes were used for normalization (n=4 independent experiments, P= 0.0018, <0.0001, 0.0011). **L.** Import of synthetic miR-146a and miR-155 into endosomes. Synthetic miR-146a or miR-155 (10nM) was incubated and, after the 30 min of import assay and RNases treatment of endosomes, the imported miRNA was recovered and quantified by qRT-PCR and value at 0 min of incubation was considered as unit. (n= 3 independent experiments, P= 0.0039 for miR-146a and P= 0.4496 for miR-155). **M.** Differential packaging of miRNAs into endosomes from a small RNA pool. Small RNA pool from C6 cells were incubated with isolated endosomes and level of imported and RNase protected level of different miRNAs were measured. The imported miRNA was quantified by qRT-PCR and in each case, value without small RNA pool added in incubation was considered as unit. Endogenous miR-146a level present in endosomes were used for normalization (n≥3 independent experiments, P= 0.0035, 0.0004, 0.0531, 0.0073 and 0.0009). Data information: In all the experimental data, error bars are represented as mean with SD, ns, nonsignificant, *P < 0.05, **P < 0.01, ***P < 0.001, ****P < 0.0001, respectively. P-values were calculated by two-tailed paired t-test in most ofthe experiments unless mentioned otherwise. Relative level of miR-122 was normalized with miR-146a level by 2^-ΔΔCt^ method. Fold change of miRNA were calculated by 2^-ΔCt^ method. Positions of molecular weight markers are marked and shown with the respective Western blots.

We have used synthetic single stranded miRNA as import substrate. However, usually miRNAs found complexed with Ago proteins in mammalian cells and Ago2-miRNA RISC complex is the functional entity for miRNA-mediated translation repression *in vivo*. Does miRNA complexed with Ago2 protein be the substrate of import assay we developed? Purified Ago2 miRNPs were incubated with isolated endosomes and RNase protected miRNA content was measured after the reaction. Like the observation we had for synthetic miR-122, the use of miR-122-Ago2 complex as substrate also showed a time dependent entry of miR-122 inside the endosomes that were protected from RNase (Fig S2).

**Figure 2.**
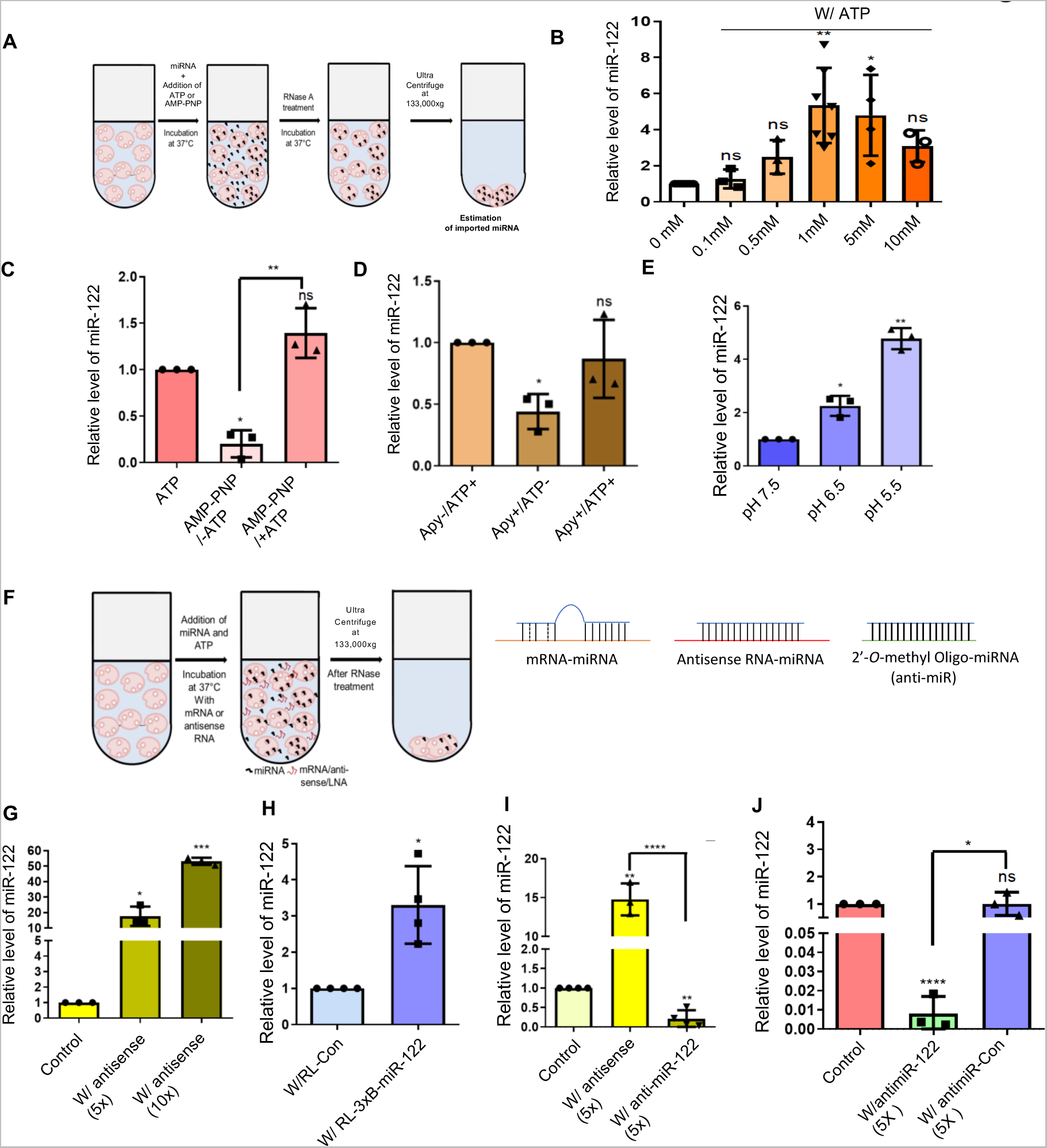
Import of miRNA to endosome is a ATP dependent process. **A.** Schematic diagram of the experimental setup to determine the energy requirement of the import process **B.** Effect of ATP concentration in the reaction buffer on miRNA packaging into endosomes in the cell-free assay (n≥3 independent experiments, P= 0.466, 0.1081, 0.0015, 0.043 and 0.0549). Import level at 0 mM ATP considered as unit. Relative levels of imported and RNase protected miRNA was measured by qRT-PCR and plotted. **C.** Effect of AMP-PNP (non-hydolyzable analogue of ATP) on miRNA import into endosomes in presence and absence of ATP (n=3 independent experiments, P= 0.0109, 0.1256 and unpaired t test P= 0.0025). Relative levels of imported and RNase protected miRNA was measured by qRT-PCR and plotted. Import level in ATP supplemented complete reaction condition was considered as unit. **D.** Effect of depletion of ATP by Apyrase and ATP supplementation after apyrase treatment on miRNA import process into endosomes (n=3 independent experiments, P= 0.0208, 0.5443). Relative levels of imported and RNase protected RNA was measured by qRT-PCR and plotted. Import level in ATP supplemented complete reaction condition was considered as unit**. E.** Effect of variation in pH of the reaction buffer on incorporation of synthetic miR-122 into endosomes. (n=3 independent experiments, P= 0.0289, 0.0037). Import level in ATP supplemented complete reaction condition at pH 7.5 was considered as unit. Relative levels of imported and RNase protected RNA was measured using qRT-PCR and plotted. **F.** Schematic diagram of experimental setup to test the effect different antisense oligonucleotides on miR-122 import. Possible hybridized structures of miR-122 with oligonucleotides have been depicted. **G.** Effect of antisense sequence containing RNA on miR122 import. Levels of imported miR-122 in absence and presence of anti-miR-122 RNA has been estimated by qRT-PCR and relative levels of imported and RNase protected RNA were plotted. Level of import in absence of inhibitory oligos was consider as unit (n=3 independent experiments, P= 0.0439, 0.0007). **H. E**ffect of mRNA having three miR-122 binding sites on miR-122 import in the *in vitro* reaction. Import level of miR-122 in presence of RL-con, an mRNA without miR-122 binding sites, was considered as unit (n=4 independent experiments, P= 0.0231). **I.** Anti-miR-122 inhibitory oligos (2’-O-Me-anti-miR122) caused drastic downregulation of miRNA packaging. Levels of imported miR-122 in absence and presence of anti-miR-122 RNA or 2’-*O*-Me-anti-miR122 have been estimated by qRT-PCR and relative levels of imported and RNase protected RNA were plotted. Level of import in absence of inhibitory oligos was consider as unit (n≥3 independent experiments, P= 0.0073, 0.0055, unpaired t-test P < 0.0001). **J.** Non-miR-122 targeting 2’-*O*-Me-miR-control have no inhibitory effect on miR-122packaging. Levels of imported miR-122 in absence and presence of 2’-*O*-Me-anti-miR122 or 2’-*O*-Me anti-miR-control were estimated by qRT-PCR and relative levels of imported and RNase protected RNA were plotted. Level of import in absence of inhibitory oligos was consider as unit (n=3 independent experiments, P= <0.0001, 0.986, unpaired t-test P= 0.015). Data information: In all the experimental data, error bars are represented as mean with SD, ns, nonsignificant, *P < 0.05, **P < 0.01, ***P < 0.001, ****P < 0.0001, respectively. P-values were calculated by two-tailed paired t-test in most of the experiments unless mentioned otherwise. Relative level of miR-122 was normalized with miR-146a level by 2^-ΔΔCt^ method.

### Requirement of ATP hydrolysis and pH change for endosomal entry of miRNA

The trafficking of charged molecules, like RNA, across the biological membrane are usually energy dependent processes. Import of tRNA through the mitochondrial double membrane or mRNA export from nucleus to cytoplasm require ATP hydrolysis (Bhattacharyya and Adhya, 2004; Xie and Ren, 2019).We wanted to check the requirement of different biochemical factors including ATP for the endosomal import of miRNAs. To understand the energy dependency of this cell-free reaction, miRNA import assays were performed in presence or absence of ATP (Fig 2A). With increasing concentration of ATP added in the *in vitro* assay, presence of 1mM ATP was found to be optimal for packaging of miRNA into endosomes (Fig 2B). However, the isolated endosomes are not free from residual ATP to cause a basal level of miRNA impot happening in absence of externally added ATP. Thus, the exclusive requirement of ATP in this process was required to be verified. In subsequent experiment, we lowered the level of ATP in isolated endosomes by pre-treating them with ATP utilizing enzyme Apyrase to deplete the endogenous ATP pool associated with isolated endosomes. In other set of assays, we have used a non-hydrolysable analogue of ATP (AMP-PNP) in the reaction buffer to compete out endogenous ATP. In both cases, depletion of ATP or use of AMP-PNP lowered the amount of packaged miRNA levels. Interestingly, exogenously added ATP could restore the miRNA import in apyrase-treated endosomes (Fig 2C, D). These data reconfirmed the importance of ATP in miRNA import process where hydrolysis of ATP provides the energy for trafficking of miRNA into endosomal lumen.

During endosome maturation, pH of the lumen of the vesicle becomes acidic as the early endosomes matures to MVBs. Does this change of pH of the reaction buffer influences the miRNA packaging process? With isolated endosome resuspended in reaction buffers of various pH, in which the *in vitro* reactions were carried out, we observed, higher amount of miRNA packaged in endosomes incubated in acidic pH compared to in endosomes incubated at pH 7.5. Thus, the acidic pH favours the endosomal packaging of miRNAs (Fig 2E).

### Target mRNA-uncoupling is required for endosomal miRNA targeting

In which form the miRNAs (single or double stranded) is best suited for getting imported inside the endosomes? Does the double stranded or a mRNA bound partial double stranded form of miRNA have restricted importability to endosomes? We have performed the cell-free reaction with isolated endosomes with miR-122 and have checked if the presence of mRNA with miR-122 binding sites or complementary anti-sense oligonucleotides has any effect on the miR-122 packaging process to endosomes. For that, we pre-incubated synthetic miR-122 with *in vitro* transcribed mRNA, or synthetic antisense RNA, or 2’-*o*-methyl modified anti-miR-122 oligo and then added the preformed miRNA-oligo/mRNA hybrids to the *in vitro* reaction with isolated endosomes and followed the miRNA entry (Fig 2F). When, we pre-incubated the synthetic miR-122 (22 nucleotides) with a 36-nucleotide long RNA, containing antisense sequence of miR-122, we found a positive effect of antisense miR-122 RNA on packaging of miR-122 (Fig 2G). We synthesized full length m7G-capped and polyadenylated mRNAs with or without three miR-122 imperfect binding sites (RL-Con and RlL-3xB-miR-122) and synthetic miR-122 was pre-incubated with RL-Con or RL-3xB-miR-122 mRNA. We observed an upregulated import of miR-122 in presence of RL-3xB-miR-122 mRNA compared to RL-con. This data indicates a positive effect of mRNA having miR-122 binding sites on cognate miR-122 import (Fig 2H). Similar experiments were carried out in presence of anti-miR-122 oligos (2’-O-methylated RNA oligos). The 2’-O-methylated anti-sense RNA oligos have strong binding affinity for sense miRNA than that of non-modified antisense RNA oligos to form stable double stranded hybrid with cognate miRNA in solution. Compared to non-modified RNA antisense oligos the 2’-*O*-methylated anti-miR-122 RNA oligos found to have an inhibitory effect on miR-122 import, suggesting the stable double stranded form of miR-122 and 2’-*O*-methylated anti-miR-122 may not be the suitable substrate for import. The chemical nature of 2’-*O*-methylated RNA did not have impact on miRNA import as the anti-miR-control 2’-*O*-methyl RNA with same modification but without the miR-122 complementarity had no inhibitory effect on miR-122 import process (Fig 2J). However, the weak double stranded form of miRNA-mRNA hybrid may get recognized by the import machinery more efficiently than the single stranded form to cause an enhanced miRNA import noted with partial or week double stranded form of substrate miRNA. However, the weak double stranded miRNA-mRNA hybrid subsequently may get uncoupled to yield single stranded miRNA for the import process to complete (Fig 2I).

### HuR is necessary for miRNA import to endosomes

Human ELAV protein HuR plays a key role in exporting miR-122 via exosomes under metabolic stress in hepatocytes. Role of HuR in export of other miRNAs have also been observed in macrophage and non-macrophage cells as well (Goswami et al., 2020; Mukherjee et al., 2016). HuR binding-unbinding to target miRNAs found to be important for miRNA export to occur (Mukherjee et al., 2016). How does HuR contribute to miRNA loading into endosomes for subsequent export of miRNAs? Although, HuR is predominantly localized in nucleus, it does shuttle to cytoplasm under specific conditions like cellular stress. As observed in stressed hepatic cells, we did find a fraction of HuR protein to be localized with the endosomal fractions isolated from C6 cells (Fig 3A). To validate that HuR has a role in miRNA import process in the cell-free *in vitro* assay system, we used purified recombinant HuR (rHuR) protein that can bind miR-122 (Mukherjee et al., 2016). We pre-incubated variable amount of the recombinant HuR protein with synthetic miR-122 to carry out the cell-free miRNA import assay with isolated endosomes (Fig 3B).The recombinant HuR positively influences the packaging of synthetic miR-122 in a concentration dependent manner, although the positive effect of HuR gets diminished at a higher concentration of HuR (Fig 3C).In an alternative approach, we overexpressed HA-HuR in C6 cells and then isolated endosomes from these cells along with control set of endosomes prepared from non-HA-HuR expressing cells to perform the *in vitro* miRNA import assay. The endosomes from HuR overexpressed cells showed several folds enhanced internalized synthetic miR-122 than the control set of endosomes under identical reaction conditions. In that experiment, we also used a deletion mutant of HuR (HuRΔH), lacking the hinge region present within two RNA recognition motifs (RRMII and RRMIII). Deletion of the hinge region makes HuRΔH unable to get uncoupled with the bound miRNA reversibly as it failed to get ubiquitinated -a process necessary for HuR-miRNA unbinding and miRNA export to occur (Mukherjee et al., 2016). Cells overexpressing HuRΔH were taken for endosome isolation and the isolated exosomes were used to carry out the *in vitro* assay along with control and full-length HA-HuR expressing cell derived endosomes. It was observed that the endosome associated and RNase protected level of miR-122 within endosomes derived from HA-HuRΔH expressing cells was significantly lower than the full-length HA-HuR expressing sets (Fig 3F). This data confirmed the positive role of full length HuR in endosomal miRNA import. miRNA unbinding of HuR is possibly required for miRNA loading to endosomes in vitro. In similar experimental context, using recombinant HuR in the *in vitro* reaction or using endosomes from HA-HuR expressing cells, we documented enhanced import of synthetic miR-146a in presence of HuR (Fig 3D, G). To confirm the role of HuR protein in the endosomal miRNA import process, we inhibited the activity of HuR by blocking the protein with antibodies specific for HuR. We observed lowered miR-122 packaging when HuR was blocked with α-HuR antibody (Fig 3E) compared to control IgG treated set.

**Figure 3.**
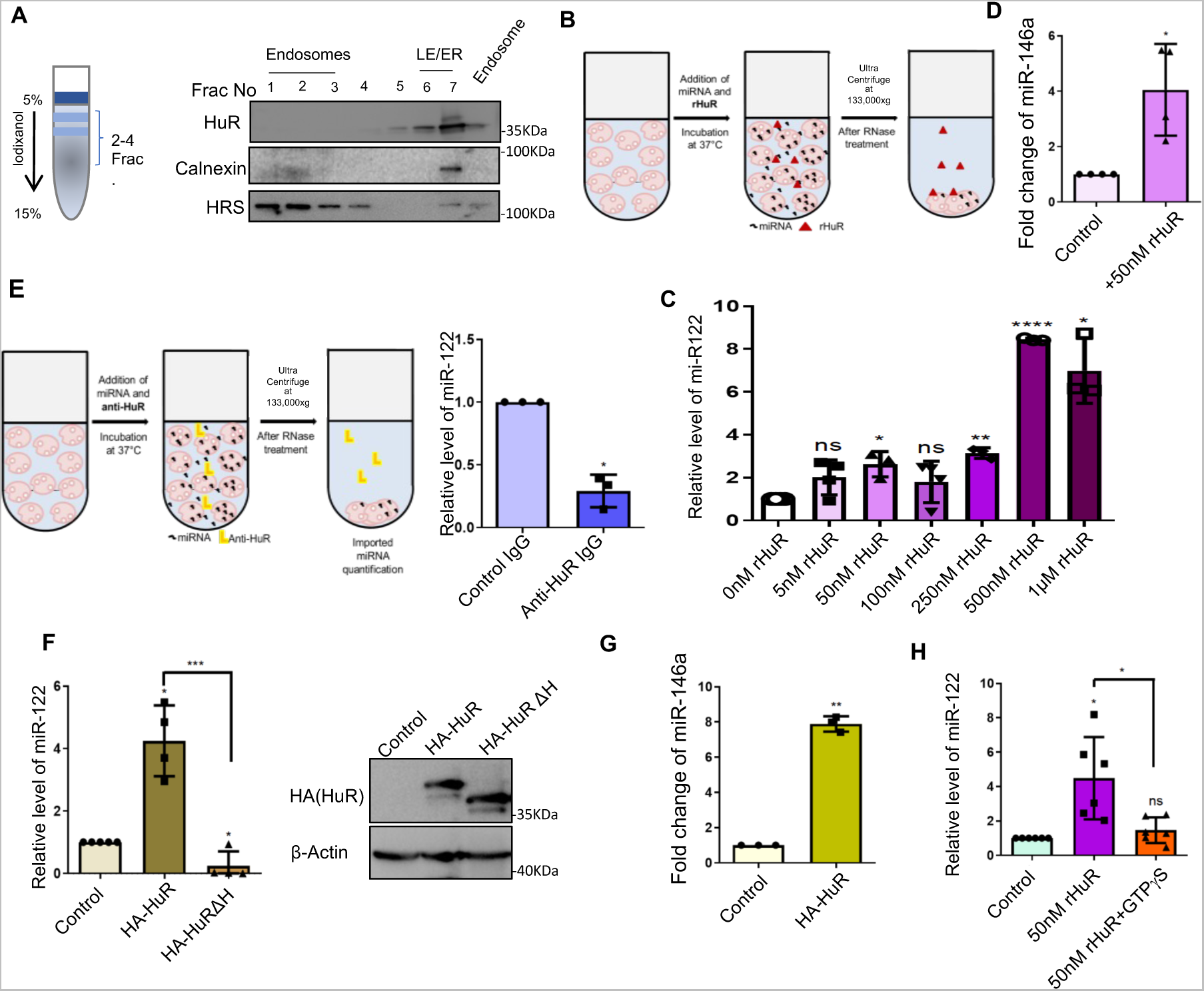
miRNA binder protein HuR is necessary for endosomal import of miR-122. **A.** Western blot analysis of the fractions of C6 cell lysate from a 5-15% Optiprep gradient for different proteins. Endosomes recovered from 1-3^rd^ fractions were also analysed (right most lane) **B.** Schematic diagram of cell-free endosomal import reaction of miR-122 performed in presence of purified recombinant HuR (rHuR)**. C.** Effect of recombinant HuR (rHuR), used at different concentrations, on packaging of synthetic miR-122 into endosomes. Import levels of miR-122 were estimated by qRT-PCR and relative levels of imported and RNase protected RNA were plotted. Level of miR-122 in absence of HuR was consider as unit (n≥3 independent experiments, P= 0.0882, 0.0414, 0.1995, 0.0044, <0.0001, 0.0206 and 0.2081). **D.** Effect of recombinant HuR (rHuR ; 50nM), on packaging of synthetic miR-146a into endosomes. Import levels of miR-146a were estimated by qRT-PCR and relative levels of imported and RNase protected RNA were plotted. Level of miR-146a imported in absence of HuR was consider as unit (n=3 independent experiments, P= 0.0348). **E.** Schematic diagram of cell-free reaction performed to test effect on anti-HuR antibody on miR-122 import (left panel). The level of miR-122 imported in presence of anti-HuR antibody was compared to control IgG treated set. Import levels of miR-122 were estimated by qRT-PCR and relative levels of imported and RNase protected RNA were plotted. Level of imported miR-122 in presence of control IgG was consider as unit (right panel; n=3 independent experiments, P= 0.0111). **F.** Effect of deletion of the hinge region of HuR, necessary for its ubiquitination and miRNA unbinding, on endosomal miRNA import. Full length HuR (HA-HuR) and its mutant version with deletion in hinge region (HA-HuRΔH) were expressed in C6 cells and endosomes were isolated. In vitro miRNA import reaction was performed using synthetic miR-122 as substrate and internalized miR-122 level was measured by qRT-PCR and relative levels of imported and RNase protected RNA were plotted. Level of miR-122 imported in HA-HuR expressing cell derived endosome set was consider as unit (n=4 independent experiments, P= 0.0106, 0.0467, unpaired t-test P= 0.0006). Immunoblots indicate expression level of HA-HuR and HA-HuRΔH in C6 cells used for endosome isolation. **G.** Cell-free reaction with endosomes from HA-HuR expressing cells showed positive effect of HuR on synthetic miR-146a packaging also. Internalized miR-146a level was measured by qRT-PCR and relative levels of imported and RNase protected miRNA were plotted. Level of miR-146 imported in the non-HA-HuR expressing cell derived endosome set was consider as unit (n=3 independent experiments, P= 0.0013). **H.** Non–hydrolyzable GTP analogue (GTPγS) negatively affects miRNA packaging into endosomes even in presence of rHuR. Cell-free in vitro import reaction with recombinant HuR was carried out in presence or absence of GTPγS were carried out and internalized miR-122 level was measured by qRT-PCR and relative levels of imported and RNase protected RNA were plotted. Level of miR-122 imported in absence of rHuR and GTPγS was considered as unit (n=6 independent experiments, P= 0.016, 0.184, unpaired t test P= 0.0145). Data information: In all the experimental data, error bars are represented as mean with SD, ns, nonsignificant, *P < 0.05, **P < 0.01, ***P < 0.001, ****P < 0.0001, respectively. P-values were calculated by two-tailed paired t-test in most of the experiments unless mentioned otherwise. Relative level of miR-122 was normalized with miR-146a level by 2^-ΔΔCt^ method. Fold change of miRNA were calculated by 2^-ΔCt^method. Positions of molecular weight markers are marked and shown with the respective Western blots.

### RalA GTPase is required for endosomal entry of miRNAs

How does HuR help in the entry of miRNAs into endosomes? Interestingly, the entry of miRNA into endosomes got inhibited by non-hydrolysable analogue of GTP (GTPyS) present in the reaction buffer (Figure 3H). There are several GTPase proteins associated with endosomes, such as Rab family of GTPases. Few of the ESCRT class of proteins or Ras superfamily of small GTPases are also present in endosomes (Bucci et al., 1992; Lebrand et al., 2002; Raiborg and Stenmark, 2009). They can possibly use the GTP hydrolysis to accelerate the entry of miRNAs across the endosomal membrane. We have used an increasing concentration of GTP in the *in vitro* miR-122 packaging reaction and observed that even 1mM GTP could enhance miR-122 packaging significantly. The optimal concentration of GTP in the import assay was found to be 5mM. However, a further increase of GTP concentration negatively affected miRNA packaging (Fig 4A). Next, to compete out the pre-existing endogenous GTP, we used a non-hydrolyzable analog of GTP (GTPyS), to observe its effect on endosomal miRNA packaging. The import of miRNA gets substantially affected in presence of GTPyS. When GTP was supplemented to the GTPyS containing reaction, the miRNA packaging could be partially rescued (Fig 4B). From these findings we concluded that, like ATP, miRNA packaging is dependent also on GTP hydrolysis, which signifies the requirement of specific GTPase protein in miRNA import reaction. To look for a candidate GTPase protein that may work in conjunction with HuR to package miRNA into endosomes, we tried to identify a GTPase protein that has interaction or association with the protein HuR. As the import process is HuR dependent, we performed HuR immunoprecipitation assays from C6 cellular lysates to get the GTPase associated with it. From the immunoprecipitation reaction, one small GTPase protein RalA (Hyenne et al., 2015), one of the two Ral proteins that belongs to the Ras superfamily of small GTPase proteins, was found to be associated with HuR (Fig 4C, D). Does RalA have a role in miRNA export? We analysed the miRNA content of extracellular vesicles (EVs) released by FH-RalA expressing C6 cells and have observed an increased quantity of miR-122, miR-146a and let-7a in the EVs secreted FH-RalA expressing cells (Fig 4E). Moreover, by NTA study of EVs, it was also observed that due to FH-RalA expression, the number of released EVs also increased significantly possibly due to increased biogenesis and packaging process happening at multivesicular endosomes in cells expressing FH-RalA (Fig S3A and B). We analysed the RalA subcellular distribution in C6 cells and have noted association of RalA with the isolated endosomes. STX5 is a SNARE protein that is known to promote fusion of multivesicular endosomes with plasma membrane to release EVs out of the cell (Linders et al., 2019). We detected both RalA and STX5 proteins in the endosomal fractions, but relative level of RalA found to be higher in isolated endosomes than STX5 (Fig 4G). We further studied the role of RalA and STX5 in miRNA import using the *in vitro* assay system. We used endosomes isolated from FH-RalA or FH-STX5 expressing C6 cells along with endosomes from a control set. There was higher packaging of miR-122 in endosomes from FH-RalA expressing cells than that in endosomes from control or FH-STX5 expressing cells (Fig 4F).Next, to further confirm the importance of RalA in miRNA import across the endosome membrane, we inhibited the activity of RalA by using α-RalA antibody in the *in vitro* assay system. Hindrance of RalA activity by α-RalA antibody, miR-122 packaging was found to be retarded significantly (Fig 4H, left panel).

**Figure 4.**
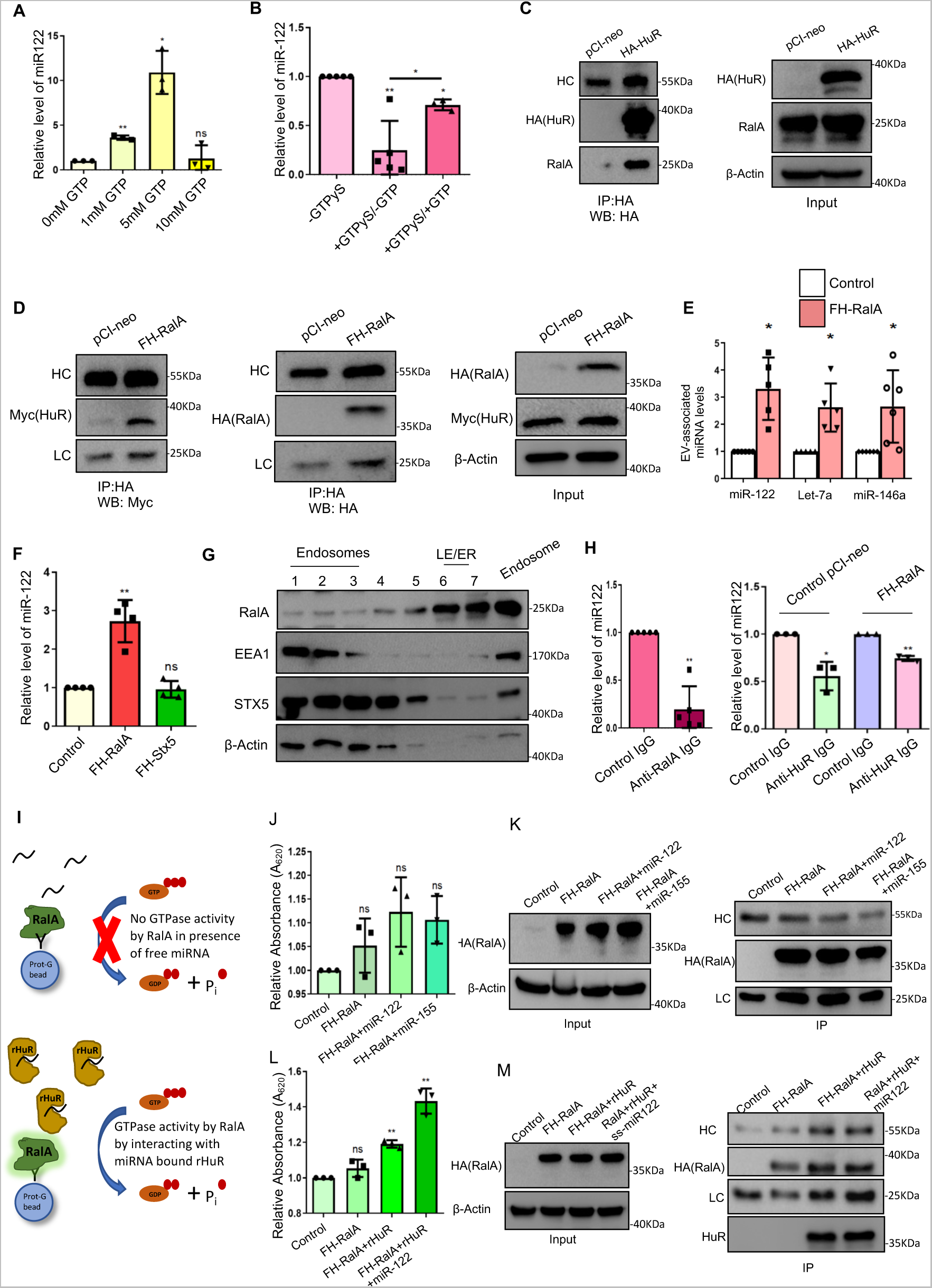
RalA GTPase drives miRNA packaging to endosomes by interacting with HuR. **A**. Effect of GTP on miRNA packaging in cell-free reaction with isolated endosomes. Internalized miR-122 level was measured by qRT-PCR and relative levels of imported and RNase protected RNA were plotted. Level of miR-122 imported in reaction with 0 mM GTP was considered as unit (n=3 independent experiments, P= 0.003, 0.0192, 0.7825) **B.** Decrease in miRNA packaging when non–hydrolyzable GTP analogue (GTPγS) was used, which was rescued by supplementary GTP. miRNA import reactions were carried out in presence and absence of GTP or GTPγS and internalized miR-122 level was measured by qRT-PCR and relative levels of imported and RNase protected miRNA were plotted. Level of miR-122 imported in absence of GTP or GTPγS was consider as unit (n≥3 independent experiments, P= 0.0049, 0.0121, unpaired t-test P= 0.0421) **C.** Western blot of RalA and HA-HuR done with immunoprecipitated materials obtained with anti-HA antibody from cell lysates of C6 cells expressing HA-HuR. PCI-neo control plasmid transfected C6 cells lysate not expressing HA-HuR was used as control. HA-HuR and endogenous RalA were detected with HA and RalA specific antibodies. Input from total lysate showing expression of HA-HuR and RalA in control and HA-HuR expressing cells. **D.** Western blot for FH-RalA and HuR in FH-RalA immunoprecipitated material obtained with anti-HA antibody from Myc-HuR expressing cell lysates also expressing or not expressing FH-RalA. Input from total lysate showing expression of Myc-HuR and FH-RalA in control and FH-RalA expressing cells also expressing Myc-HuR. **E.** miRNA content of EVs isolated from control and FH-RalA expressing cells. Control cells were transfected with pCI-neo vector. miRNA levels were measured by qRT-PCR. Levels of miRNA in EVs from control cells not expressing FH-RalA imported were consider as units ((n≥5 independent experiments, P= 0.0108, 0.0149, 0.0279). **F.** Levels of miR-122 imported to isolated endosomes from control and FH-RalA or FH-Stx5 expressing cells. miR-122 level was measured by qRT-PCR and relative levels of imported and RNase protected RNA were plotted. Level of miR-122 imported in the pCI-neo transfected control cell derived endosome was consider as unit (n≥3 independent experiments, P= 0.0081, 0.7197, 0.4621). **G.** Western blot of subcellular fractionations of C6 cell lysate separated on 5-15% Optiprep gradient for detection of RalA and STX5. Fractions enriched for early endosomes were pooled to get endosome enriched fraction and was also analyzed by western blot (right most lane). **H.** The level of miR-122 imported in presence of anti-RalA antibody was compared to control IgG treated set. Import levels of miR-122 were estimated by qRT-PCR and relative levels of imported and RNase protected RNA were plotted. Level of miR-122 in presence of control IgG was consider as unit (n=5 independent experiments, P= 0.0017, left panel). In right panel effect of anti HuR antibody on FH-RalA driven import of miRNA is shown. The level of miR-122 imported in presence of anti-HuR antibody was compared to control IgG treated set both for pCI-neo and FH-RalA expressing cell derived endosomes. Import levels of miR-122 were estimated by qRT-PCR and relative levels of imported and RNase protected RNA were plotted. Level of miR-122 in presence of control IgG was consider as unit. **I.** Diagram of RalA activity assay measurement in presence and absence of miRNA and HuR. FH-RalA expressed in C6 cells was immobilized on FLAG-beads. Immobilized FH-RalA GTPase hydrolyzes GTP in the reaction and free inorganic phosphate level was measured by malachite green based detection system. **J.** Presence of synthetic miRNA alone does not alter GTPase activity of RalA significantly. GTPase assay was done in absence and presence of miRNA and released P_i_ levels were measured and relative activity was plotted. Activity with bead control was used as unit. (n=3 independent experiments, P= 0.2562, 0.1007, 0.0669). **K.** Western blots of FLAG-bead immobilized FH-RalA and input from total cell lysate used for immunoprecipitation. **L.** Presence of rHuR in the reaction increased RalA activity which was further increased when rHuR was bound to miR-122 was added in the reaction. GTPase assay was done in presence and absence of recombinant HuR (50nM) in presence of synthetic miR-122 (10nM) and released P_i_ levels were measured and relative activity was plotted. Activity with bead control was used as unit. (n=3 independent experiments, P= 0.2, 0.004, 0.0089). **M.** Western blots of FLAG-bead immobilized FH-RalA and input from of total cell lysate used for immunoprecipitation in panel L. Data information: In all the experimental data, error bars are represented as mean with SD, ns, nonsignificant, *P < 0.05, **P < 0.01, respectively. P-values were calculated by two-tailed paired t-test in most of the experiments unless mentioned otherwise. Relative level of miR-122 was normalized with miR-146a level by 2^-ΔΔCt^ method. Exosomal miRNA levels were calculated by 2^-ΔCt^ method. Positions of molecular weight markers are marked and shown with the respective Western blots. HC; heavy Chain

### HuR-miRNA complex activates the GTPase RalA for miRNA targeting to endosomes

How does HuR and RalA cooperate in the process of miRNA import? Is RalA a mediator of the HuR-driven miRNA packaging into endosomes? We performed GTPase activity analysis *in vitro* for the RalA from isolated endosomes. As RalA is a GTPase protein, analysis of GTP hydrolysis and measurement of free inorganic phosphate (P_i_) release due to the GTPase activity could indicate the activity of RalA. We immobilized FH-RalA on agarose beads and carried out a reaction in presence of GTP. We incubated the RalA-immobilized beads along with GTP and either with free miRNA, recombinant HuR alone or miRNA-recombinant HuR complex. After the reaction, we measured free P_i_ level in the supernatant by a malachite green based colorimetric assay (Fig 4I). The RalA activity increased in presence of HuR which was further increased with miRNA-bound HuR, although presence of free miR-122 or miR-155 failed to generate any significant increase in GTPase activity of RalA (Fig J, L). FH-RalA levels in recovered FH-RalA containing beads were measured by western blot analysis (Fig 4K, M). The activity measurement of RalA, suggests RalA as the downstream effector of HuR, which upon interaction with HuR-miRNA complex, utilizes the GTP to package miRNAs into endosomes.This data also suggests that HuR works upstream of RalA-mediated endosomal miRNA entry. Interestingly, we could inhibit miRNA import into endosomes isolated both from control or FH-RalA expressing cells with anti-HuR antibody *in vitro*. We noted in both cases presence of anti-HuR antibody reduces the importability of miRNAs into endosomes that signifies role of HuR in miRNA import upstream of RalA (Figure 4H, right panel).

### *In vitro* Maturation of endosomes requires ATP

RalA GTPase is known to play a part in endosome maturation process and thus believed to contribute in a positive manner to form MVBs from endosomes in the *in vivo* context to facilitate the miRNA incorporation into MVBs and their export. We have observed increased number and size of EVs released by FH-RalA expressing cells compared to EVs isolated from control group of cells (Figure S3).

While exploring the various factors for their roles on miRNA packaging into endosomes in this cell-free *in vitro* assay system, we were interested to visualize the endosomes and the extent of their maturation during the *in vitro* reaction. For the visual analysis of these organelles out of their natural environment, we used AFM and NTA for direct imaging and biophysical characterization of these vesicles. We have carried out an *in vitro* assay for 30 minutes and reisolated the vesicles, resuspended them and processed for AFM or NTA analysis (Fig 5A). Along with these, we have also analyzed the endosomes after 0 min of incubation or right after isolation before the *in vitro* assay. From the atomic force micrographs and particle analysis, it was observed that pre-assay vesicles and re-isolated vesicles at 0 min post-assay showed similar morphology and size distribution (Fig 5B, C), but the 30 mins post-assay vesicles showed bigger vesicles with embedded smaller vesicles in the AFM images while a distinct peak for larger vesicles noted in particle tracking analysis (Fig 5D). These morphological analyses suggest endosomal maturation happening in this minimal in this *in vitro* system during the assays.

**Figure 5.**
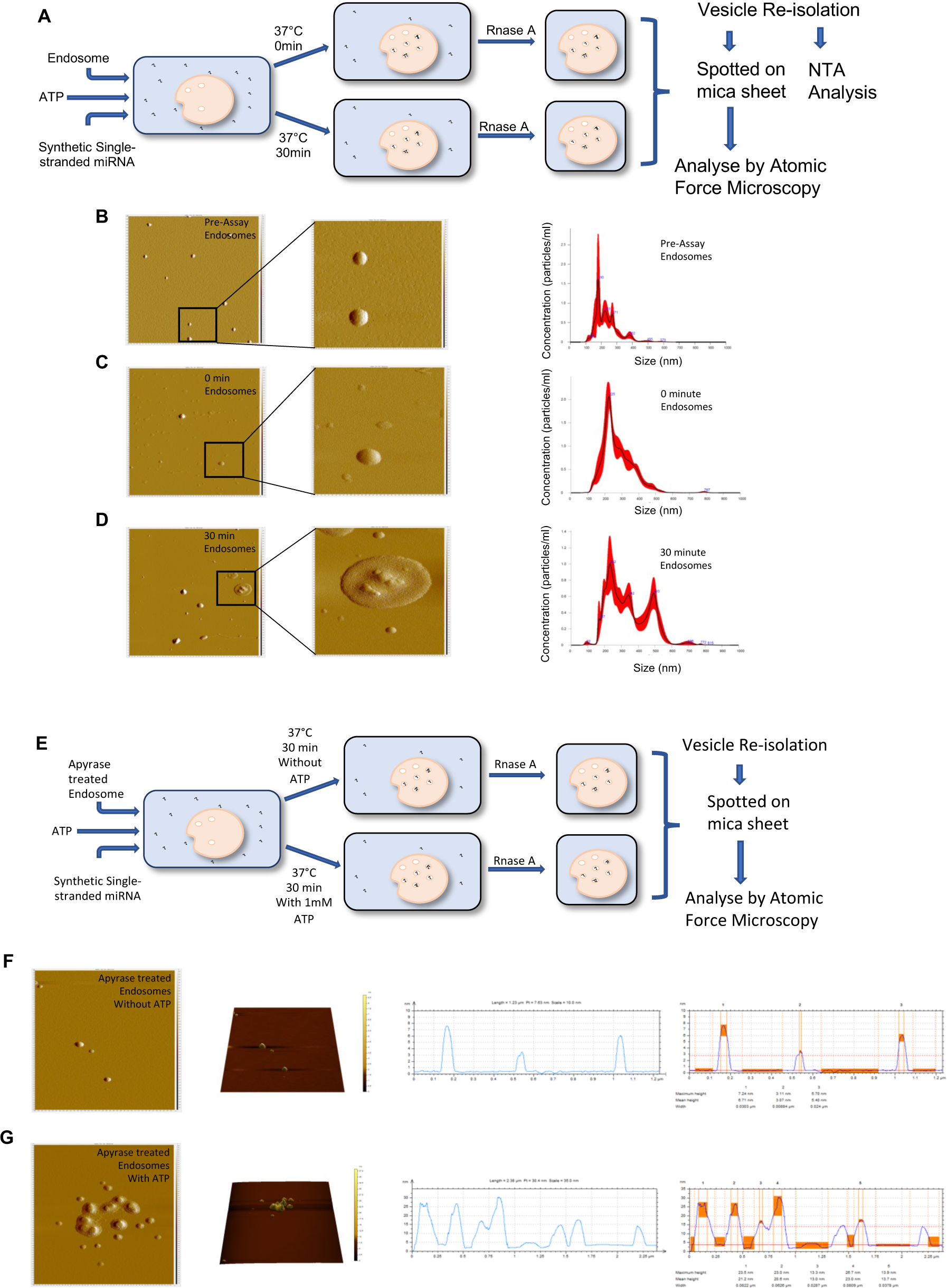
Isolated endosomes maturates to multi-lamellar structure containing organelles *in vitro* in an ATP dependent manner. **A** Schematic diagram of the study of *in vitro* maturation of isolated endosomes. **B-D.** Atomic Force micrograph of isolated endosomes prior to cell-free assays (B); or after 0 min (C) or after 30 mins (D) of incubation. Insets show zoomed higher resolution images. Analysis of their concentration and size by nanoparticle tracking Analysis (NTA). **E.** Schematic diagram of the study of *in vitro* maturation of isolated endosomes after Apyrase treatment**. F-G.** Atomic force micrograph of isolated endosomes treated either with ATP depleting enzyme Apyrase alone prior to cell-free assay (F) or supplemented with ATP after the Apyrase treatment (G). Side panels show 3D topography and amplitude graphs indicating size of vesicles.

The energy dependency of the miRNA packaging process may be linked to endosome maturation. We noted in previous experiments that ATP hydrolysis is essential for miRNA import. We were interested to check if the endosomal maturation gets affected by ATP. We collected the endosome-enriched fractions from the sub-cellular fractionation of C6 cells and treated them with ATP depleting enzyme Apyrase before the Apyrase treated endosomes were re-isolated and used for in vitro endosome maturation studies. We carried out an *in vitro* reaction in presence or absence of ATP and reisolated vesicles were analyzed by AFM imaging (Fig 5E). The AFM images of Apyrase treated endosomes showed small vesicles sporadically present over the field of view, but the ATP re-supplemented vesicles showed clustering of vesicles together and supposedly in the process of fusion and formation of larger vesicles due to presence of ATP possibly due to activity of various ATPase present on the endosomal surface (Fig 5F, G). Together, these visual analyses of endosomes suggest an active energy-dependent maturation process, capable of occurring in the *in vitro* system, and by this maturation process, miRNA as cargo may get actively packaged within endosomes.

### Rab5-mediated endosomal maturation is required for miRNA export via EVs

Studying the maturation process of endosomes made us to think about how miRNA packaging, localization and export will be affected if the endosomal maturation pathway is blocked. We used wild type and couple of mutant variants of the early endosomal protein Rab5A to explore its role in endosome maturation and miRNA export. Rab5-CA is a constitutively active form of Rab5, while Rab5-DN is a dominant negative form of Rab5A (Flinn et al., 2010) . We expressed these mutants along with a fluorescent fusion construct to image endosomes (Endo-YFP) in C6 cells and visualized endosomes and Rab5-positive endosomes by indirect immunofluorescence (Fig 6A). From the observations, it was evident that Rab5-CA enhances large early endosome formation which lowered the number of total endosomes in the cells and caused a shift in the endosomal equilibrium towards early endosomes. Rab5-WT enhanced the endosomal number while Rab5-DN affects negatively the early endosome numbers in cells expressing it (Fig 6B). We also isolated endosomes from these cells and subjected them to analysis by AFM, and there also we found similar morphology of vesicles from control and Rab5-WT expressing cells, but much larger endosomes were observed from Rab5-CA expressing cells (Fig S4A-C). For the miRNA localization study, we expressed these Rab5 mutants along with pre-miR122 expressing vector in C6 cells and investigated the cellular, endosomal and the EVs-associated miR-122 content. Level of cellular miR-122 was found to be increased upon Rab5-CA expression (Fig 6D). The miR-122 level in isolated endosomes from Ra5-CA expressing cells showed miR-122 was highly enriched in early endosomes (Fig 6C). But, due to retention of miR-122 into early endosomes in the Rab5-CA context, the extracellular export by EV was highly affected and EV-associated miR-122 level found low. However, as Rab5-WT enhanced the endosome biogenesis process (Gorvel et al., 1991), the miR-122 was found to be highly enriched in EVs compared to other sets (Fig 6E). The EV analysis by NTA also showed moderate but consistent increase in number of EVs from cells expressing Rab5-WT (Fig S4D). Overall, the observations support the role of an endosomal effector protein to accelerate miRNA packaging and export by enhancing the late endosome maturation pathway. Interestingly, we had also expressed HA-HuR and the mutant HA-HuRΔH to check their effect on endosomal morphology. Wild type HA-HuR found primarily in the nucleus while the mutant HA-HuRΔH was cytoplasmic. As HuR may not be directly involved with endosome maturation process rather act as miRNA-scavenger to get them packaged into endosomes, there was no significant effect on endosome morphology or number after expressing HA-HuR or HA-HuRΔH (Fig S5). This data further signifies that the miRNA import regulation by HuR does not affect the endosome maturation directly rather acts upstream of it.

**Figure 6.**
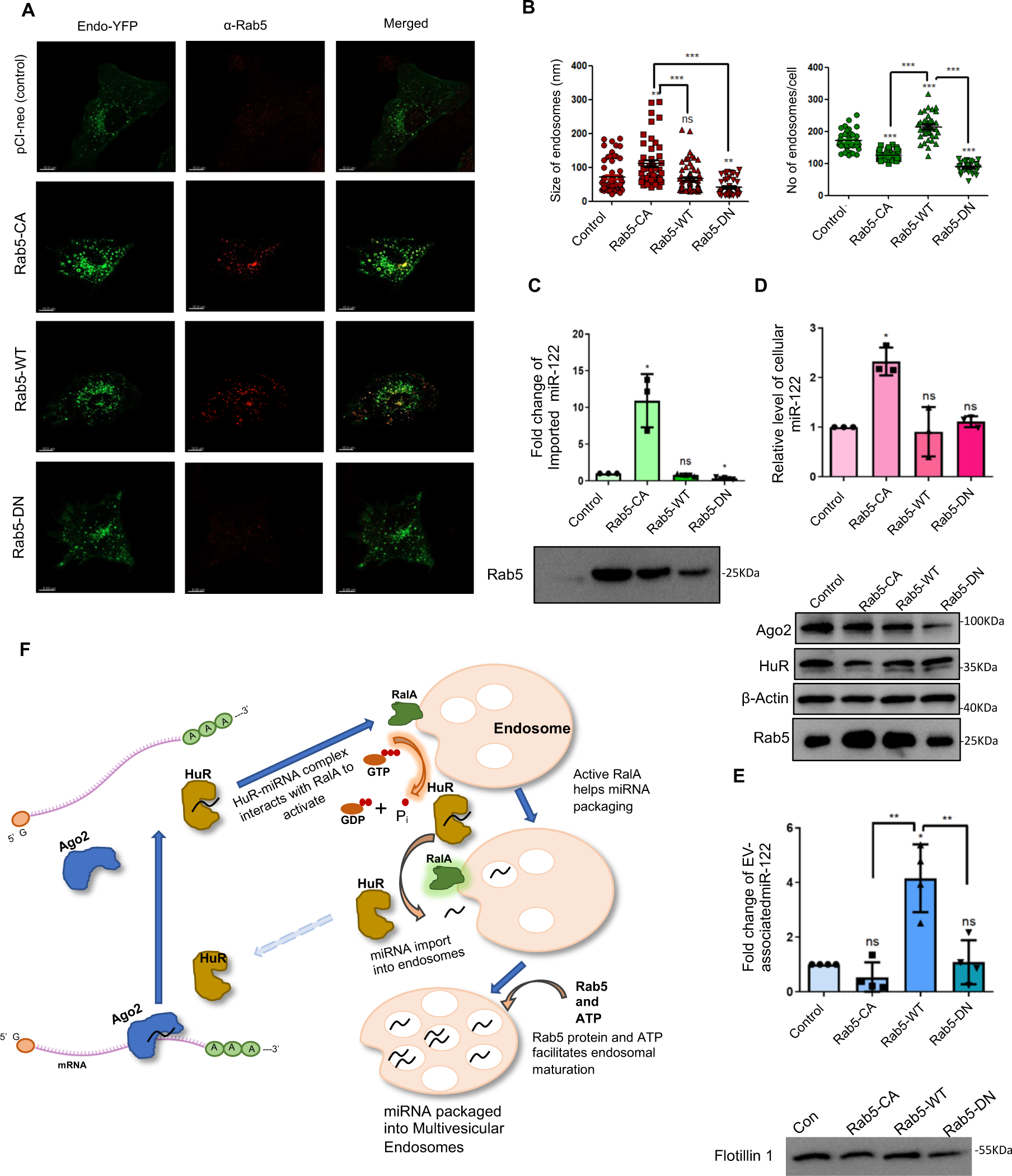
Rab5A controls endosome maturation and miRNA export. **A.** Confocal images of Endo-YFP expressing C6 cells co-expressing either of the Rab5A variants, Rab5-CA (Constitutively active), Rab5-WT (Wild type) and Rab5-DN (Dominant negative). Control set expressing pCI-neo plasmid along with Endo-YFP. Endosomes were tagged by expressing Endo-YFP (green) and overexpressed Rab5A mutants were indirectly visualized by using α-Rab5 based indirect immunofluorescence. **B.** Number and size of endosomes imaged by confocal microscopy were analysed by using IMARIS software and plotted (left-side panel of endosome size-n≥40 number of fields, P= 0.0035, 0.6053, 0.0022, unpaired t-tests P <0.001; right-side panel of number of endosomes-n≥31 number of endosomes, P <0.0001 for each paired and unpaired t-tests). **C-E.** C6 cells expressing Rab5A mutants (Rab5-CA, Rab5-WT and Rab5-DN) along with miR-122 were used to prepare endosomes and EVs were isolated from cultural supernatants. Endosomal (C; n=3 independent experiments, P= 0.0417, 0.2068, 0.0191), cellular (D; n=3 independent experiments, P= 0.0147, 0.7774, 0.2266) and EV content (E; n=4 independent experiments, P= 0.1853, 0.0148, 0.8527, unpaired t-tests P= 0.0018, 0.006) of miR-122 was analysed. Levels of miR-122 were estimated by qRT-PCR and relative levels were plotted. Level of miR-122 in presence of control set transfected with pCI-neo was consider as unit (n=3 independent experiments, P= 0.0417, 0.2068, 0.0191). Immunoblot of Rab5 was done from isolated endosomes (C). Western blots of various cellular marker proteins from the cellular lysate used for endosome isolation (D). Flotilin1 level was checked from EV samples (E). **F.** A probable model of HuR-RalA cascade of miRNA packaging into endosomes. Ago2 bound miRNA associates with target mRNA. HuR binds with miRNA and dissociates miRNA from Ago2 miRNP. HuR-miRNA complex interacts with endosomal RalA inducing the GTPase activity of RalA. miRNA gets uncoupled from HuR and gets packaged into endosomes/MVB. Rab5 protein and ATP facilitates the endosomal maturation process. Data information: In all the experimental data, error bars are represented as mean with SD, ns, nonsignificant, *P < 0.05, **P < 0.01, ***P < 0.001, ****P < 0.0001, respectively. P-values were calculated by two-tailed paired t-test in most ofthe experiments unless mentioned otherwise. Relative level of miR-122 was normalized with U6 snRNA level by 2^-ΔΔCt^ method. Fold change of miRNA were calculated by 2^-ΔCt^ method. Positions of molecular weight markers are marked and shown with the respective Western blots. Scale bar 10 µm.

## Discussion

Use of cellular organelles isolated from animal cells in a cell-free reconstruction assay system is a well-accepted model to study important biological pathways where variables can be closely monitored and manipulated to reveal the mechanistic detail of the pathway investigated (Liu and Fletcher, 2009). Analysis and dissection of the endocytic pathway is a topic of interest and, till date, several new factors and their mechanism of action in the endosomal pathway are being discovered by cell biological and molecular genetic analysis. No prior report of *ex vivo* assay system development with isolated endosome was there, possibly because of limiting number of endosomes that can be recovered to do such assays. We adopted a method of enriching endosomes and verified the enrichment and characterized the isolated endosomes using nanoparticle tracking (NTA) and atomic force microscopy (AFM) before utilizing these enriched endosomal fractions in a reconstitution assay for miRNA entry. The existing assay systems for miRNA packaging in vesicles had either used synthetic lipids in specific composition to generate artificial vesicles to get the assay system ready (Kahya et al., 2001) or used crude membrane fractions with limited characterization and validation completed to qualify as authentic endosomes that are certainly not free from other major bio-membranes and organelles (Shurtleff et al., 2016). The *in vitro* system that we have used contains endosomes and factors that are required for endosomal maturation and are primarily free from ER and mitochondria “contamination” as they were separated on a density gradient and re-isolated for the assay. Structural change due to maturation of the vesicles after the *in vitro* reaction were monitored and analysed by NTA and AFM analysis.

We have closely monitored the importance of several biochemical factors in the newly developed *in vitro* reaction for miRNA packaging into endosomes and have standardized the reaction time, temperature, amount of miRNA as cargo to maintain the optimal reaction conditions and to achieve reproducibility. To nullify the false positive signal from excess reactant in the system, we have performed an RNase protection assay to select only the imported and hence RNase protected pool of RNA to analyse. We have used an endogenous and abundantly present miRNA (miR-146a in C6 cells) in endosomes for normalization of the imported miRNA-122 substrate in all the *in vitro* reaction we performed.

With this enriched endosomes and standardized miRNA packaging *in vitro* reaction, we have confirmed the energy dependence of the process by studying the effect of ATP and GTP. Previously published report from our lab suggested that the miRNA gets packaged in Extracellular vesicles as single stranded form (Ghoshal et al., 2021). Interestingly, RNA oligonucleotide comprising the anti-sense sequence of the synthetic RNA substrate facilitates the packaging of the single stranded substrate RNA strand into endosomes. Furthermore, presence of synthetic mRNA containing the target sequence of miRNA in the reaction favours packaging of miRNA. However, strong binding with 2’-O-mythlated anti-miR oligonucleotide significantly lowered packaging of cognate miRNA into endosomes. This is consistent with a previous report citing an *in vivo* study that suggests the positive role of target mRNA on miRNA export. In human cancer cells increasing target mRNA accelerate EV-mediated export of cognate “used” miRNAs (Ghosh et al., 2021).

HuR or ELAVL1 protein has been reported earlier as an exporter of miR-122 in a context-dependent manner (Mukherjee et al., 2016). Recombinant HuR (rHuR) can bind miRNA and play important role in miRNA export process. Presence of rHuR in the assay system leads to enhanced miRNA packaging into endosomes. Endosomes from HuR overexpressing cells also showed enhanced packaging of miRNA in the *in vitro* reaction. Additionally, the mutant form of HuR that lack the hinge region and thus incapable of getting unloaded from miRNA (Mukherjee et al., 2016) also fails to package miRNA in the *in vitro* reaction as well.

We had studied the requirement of GTP in the miRNA packaging assay *in vitro* and we were interested in RalA GTPase that is known for its role in vesicle trafficking (Hyenne et al., 2015). RalA was found to be present in the isolated endosomal fractions and endosomes, from cells expressing FH-RalA, successfully package miRNAs within it. HuR and RalA associate with each other. We also found HuR-miRNA complex can stimulate the GTPase activity of RalA to positively influence the miRNA import into endosomes. Previous reports suggest HuR protein has a role in RalA expression and it either upregulates RalA expression in specific cellular context or get positively associated with HuR through the long 3’ UTR of target mRNA (Lee and Mayr, 2019; Mazan-Mamczarz et al., 2008). These findings suggest a probable mechanism where miRNA exporter HuR binds to miRNA and recruits RalA for utilizing its GTPase activity to package miRNA into endosomes. Thus, we have identified the RalA GTPase to plays a major role in miRNA import into endosomes. However, we are not sure about the exact mechanism by which this GTPases utilizes the energy of GTP hydrolysis for import of miRNAs inside the endosomes. It may be coupled with a membrane pinching or recruitment of specific ESCRT components to complete the MVB formation for entrapment of miRNAs. Further study is required to answer this question.

We were interested to follow the changes in miRNA localization and export if the endosomal homeostasis is altered. We utilized a few mutants of early endosomal protein Rab5A to study the phenomenon. The Rab5A mutants, that we have used, were either constitutively active (Rab5-CA) or dominant negative (Rab5-DN). They, along with wild type Rab5 (Rab5-WT) too, were ectopically expressed in C6 cells. Confocal microscopy and atomic force microscopy validated the structural alteration of endosomes from cells expressing the Rab5A mutants. As Rab5-CA increased the early endosomal pool which possibly shifted equilibrium during the endosome maturation reaction to early endosomes, total cellular and endosomal content of miR-122, an exogenously expressed miRNA in C6 cells, was highly increased. However, Rab5-CA expressing cell derived EVs had lower miR-122 content compared to Rab5-WT. Although Rab5A may favour more endosomal packaging of miR-122, but defective endosomal maturation hindered the export process. Rab5-DN on contrary comparatively lowered the active early endosomal pool which affected both miR-122 packaging and export process.

The primary method to study miRNA packaging into endosomes should be an *in vitro* reconstitution assay with high reproducibility. These kinds of assay systems can allow us to observe molecular and cellular processes *ex vivo* where a wide number of variables can be closely monitored or controlled to follow their role in the process. On the flip side, as cellular reconstitution focuses on only a specific set of components used by cells, this does not represent the precise mechanism of the biological process, which always raises the concern on physiological nature of such assays. So, to bridge the gap between controlled subcellular re-construction and cellular physiology, the *in vitro* reconstitution approach should be complemented together with a cell-based *in vivo* approach. By studying a cellular phenomenon from both these angles can ultimately provide the detailed and complete picture. We validated the findings that were obtained from the in vitro assay system with established cell based and EV-based experiments to support the finding in parallel assays.

The assay system described in this manuscript will be a useful tool not only to study the miRNA or mRNA import into endosomes, but could be a useful system also to study membrane dynamics during the endosome maturation. It will also be an important system to further follow and study the cargo sorting on endosomes and to investigate the defects in the system in presence of factors such as amyloid proteins that are known to affect the cargo trafficking or recruitment of mTOR proteins to endosomes in neurodegeneration context (De and Bhattacharyya, 2021; De et al., 2021).

## Experimental Procedures

### Cell culture and cell transfections

HEK293 and C6 glioblastoma cells were cultured in Dulbecco’s Modified Eagle’s medium (DMEM; Gibco) supplemented with 10% heat-inactivated fetal calf serum (HI-FCS, Gibco) and 1% of Penstrep (Gibco) in a 37°C incubator in presence of 5% carbon dioxide. For HEK293 and C6 cells, all transfections of plasmids were performed using Lipofectamine 2000 reagent, (Life technologies) according to manufacturer’s protocol.

### Optiprep density gradient centrifugation

For subcellular fractionation of organelles, a 3-30% continuous gradient was prepared using Optiprep density gradient medium of 60% iodixanol solution (Sigma-Aldrich) in a buffer composed of 78mM KCl, 4mM MgCl_2_, 8.4mM CaCl_2_, 10mM EGTA, 50mM HEPES (pH 7.0). Ice cold 1X PBS was used for washing before cells were lysed using a Dounce homogenizer (Kontes glass) in a buffer constituting 0.25M sucrose, 78mM KCl, 4mM MgCl_2_, 8.4mM CaCl_2_, 10mM EGTA, 50mM HEPES (pH 7.0) supplemented with 100μg/ml of cycloheximide, 0.5mM DTT and 1X PMSF. Cellular lysate was cleared by centrifuging twice at 1,000x*g* for 5 min before overlaying gently on top of the prepared continuous gradient. Ultracentrifugation was carried out for 5 hours at 133,000x*g* on a SW60-Ti (Beckman Coulter) rotor to resolve each gradient. After centrifugation, by aspirating from top, ten fractions were collected which were processed for analysis of proteins and RNA.

### Endosome enrichment by subcellular fractionation

Optiprep density gradient medium of 60% iodixanol solution (Sigma-Aldrich) was used to prepare a 3ml 5-10-15% step gradient (1ml of lighter solution overlaid gently on 1ml of heavier gradient) in a buffer containing 78mM KCl, 4mM MgCl_2_, 8.4mM CaCl_2_, 10mM EGTA, 50mM HEPES (pH 7.0) for separation of subcellular organelles. Cells were washed with 1X PBS and homogenized with a Dounce homogenizer (Kontes glass) in a buffer containing 0.25M sucrose, 78mM KCl, 4mM MgCl_2_, 8.4mM CaCl_2_, 10mM EGTA, 50mM HEPES (pH 7.0) supplemented with 100μg/ml of Cycloheximide, 0.5mM DTT and 1X PMSF. The lysate was clarified by centrifugation at 1,000x*g* for 5 min twice and 1ml of cleared lysate layered on top of the immediately prepared 3ml gradient at 4°C. The tubes were centrifuged at 133,000x*g* for 5 hours for separation of gradient and seven fractions of equal volume were collected by aspiration from the top for endosome/MVB isolation and subsequent analysis of proteins and RNA. For Endosome enrichment, top three fractions were pooled and diluted with a diluent buffer containing 78mM KCl, 4mM MgCl_2_, 8.4mM CaCl_2_, 10mM EGTA, 50mM HEPES (pH 7.0) to fill a 4ml tube and further ultracentrifuged at 133,000x*g* for another 2 hours. After centrifugation, endosome enriched membrane pellet was resuspended in a buffer containing 0.25M sucrose, 78mM KCl, 4mM MgCl_2_, 8.4mM CaCl_2_, 10mM EGTA, 50mM HEPES (pH 7.0) supplemented with 100μg/ml of cycloheximide, 0.5mM DTT and 1X PMSF, and this endosomal suspension was kept in ice before carrying out the *in vitro* assay.

### Cell-free *in vitro* reconstitution assay

Endosome enriched suspension isolated from either HEK293 or C6 cells were divided equally for experimental sets. As per the standardization process described in the Results section, synthetic miRNA, small RNA pool, miRNP, synthetic miRNA bound with various antisense oligos were specifically used as cargo in presence of ATP or GTP and incubation was carried out for 30 minutes at 37°C, followed by an Rnase protection assay as described below and vesicles were washed and re-isolated by ultracentrifugation at 133,000x*g* for 2 hours. Vesicle pellets were processed for RNA and protein isolation and analyzed.

### RNase protection assay

To degrade excess free miRNA which were not internalized into endosomes and thus to nullify false positive signal, an RNase protection assay was carried out after *in vitro* assay, before washing and re-isolating the endosomes. DNase and protease-free RNase A (Thermo scientific) was used at the concentration of 0.5unit/µL for 5 minutes at 37°C. To ensure the effectivity of Rnase A, a reaction was carried out in presence of the detergent sodium deoxycholate to disrupt endosomal membrane and degrade even the miRNA that would have otherwise remain protected from the effect of RNase A. After RNase A reaction, to remove RNase and excess reactants, vesicles were washed by increasing the volume with diluent buffer and re-isolating them by ultracentrifugation at 133,000xg for 2h.

### Isolation of Ago2 bound miR-122 (miRNP) complex

For using Argonaute-2 bound miRNA as cargos for *in vitro* assays, HEK293 cells transiently expressing FH-Ago2 and pre-miR122 (by expression vectors) were lysed, 48h after transfection, in lysis buffer (10mM HEPES pH 7.4, 200mM KCl, 5mM MgCl_2_, 1mM DTT, 1X PMSF, 40 U/ml RNase inhibitor, 0.5% Triton X-100 and 0.5% sodium-deoxycholate) at 4°C for 30 minutes, followed by clearing of the lysate by centrifuging at 3,000x*g* for 10 mins. The cleared lysate was incubated with pre-blocked anti-FLAG M2 affinity matrix (Sigma) for immunoprecipitation for 16h at 4°C on an end-to-end rotator. IP buffer (20mM Tris-HCl pH 7.4, 150mM KCl, 5mM MgCl2, 1mM DTT, 1X PMSF) was used for washing the beads for three times at 4°C. Affinity purified FH-Ago2 bound with miRNA (most population bound with miR-122 due to its overexpression) was eluted from anti-FLAG beads by competing with 3X FLAG peptide (Sigma) as per the manufacturer’s protocol in a purification buffer (30mM HEPES pH 7.4, 100mM KCl, 5mM MgCl2, 0.5mM DTT, 3% glycerol) and isolated miRNP from the supernatant was stored at -80°C or used in the *in vitro* assay.

### Extracellular Vesicles isolation

C6 cells transfected with specific plasmids were split 24h post-transfection into 90mm plates and incubated for 24 hours to achieve 70-80% confluency. The conditioned media were subjected to EV isolation as described in previous report with minor modifications (Thery et al., 2006). Cells were grown in growth medium supplemented with EV-depleted serum to prevent any interference from exosomes or EVs present in serum. The conditioned media was clarified for cellular debris and other contaminants by centrifugation at 2,000x*g* for 10 minutes and 10,000x*g* for 30 minutes followed by filtering through a 0.22-µm filter for further clearing. EVs were isolated by ultracentrifuging the cleared conditioned media at 100,000x*g* for 90 min. When analysis of proteins was required, an additional ultracentrifugation of the cleared conditioned media was done on 30% sucrose cushion at 100,000*g* for 90 minutes, and the layer of EVs was further diluted with 1X phosphate buffered saline (PBS) by ultracentrifugation at 100,000x*g* for 90 minutes for obtaining EV pellet.

### Immunoblotting

Western blotting of proteins was performed as mentioned in previous reports (Ghosh et al., 2013; Ghosh et al., 2015). Protein samples from cellular lysate, subcellular fractionated samples or immunoprecipitated proteins were subjected to SDS-polyacrylamide gel electrophoresis followed by transfer of the same to PVDF nylon membrane overnight at 4°C. Membranes were blocked by using 3% BSA for 1 hour then specific required antibodies were used to probe the blot for at least 16h at 4°C. Membranes were incubated for 1h with horseradish peroxidase-conjugated secondary antibodies (1:8000 dilutions) at room temperature. Images of developed western blots were taken using an UVP BioImager 600 system coupled with VisionWorks Life Science software, version 6.8 (UVP).

### RNA Isolation and Real Time PCR

Total RNA was isolated by using TriZol or TriZol LS reagent (Invitrogen) according to the manufacturer’s protocol. Samples containing low amount of miRNA were precipitated by using isopropanol in presence of glycoblue co-precipitant (Thermofisher Scientific, USA). cDNA was prepared by taking 100-200ng of RNA from cellular RNA samples or by taking equal volume of RNA isolated from *in vitro* assay samples or EV samples by using a Taqman reverse transcription kit (Applied Biosystems). miRNA assays by real time PCR was performed using specific primers. Real time analyses by two-step RT-PCR were performed for quantification of miRNA levels on Bio-Rad CFX96TM real time system using Taqman (Applied Biosystems) based miRNA assay system. One third of the reverse transcription mix was subjected to PCR amplification with TaqMan Universal PCR Master Mix No Amp Erase (Applied Biosystems) and the respective TaqMan reagents for target miRNA. Samples were analyzed in triplicates. For normalization of miRNA levels of cellular samples, U6 snRNA was used, whereas for *in vitro* assay samples done with synthetic miR-122, endosomal miR-146a was used for normalizing the level of miR-122.

### Immunoprecipitation assay

For immunoprecipitation (IP) reactions, C6 cells were lysed by using lysis buffer (20 mM Tris-HCl pH 7.5, 150 mM KCl, 5 mM MgCl2 and 1 mM DTT) containing 0.5% Triton X-100, 0.5% sodium deoxycholate, 40 U/ml RNase inhibitor and 1X PMSF for 30 min at 4⁰C, followed by sonication of three pulses of 10 seconds each. Lysates were clarified by centrifuging at 16,000x*g* for 15 minutes. Protein G-agarose beads were blocked by using a solution of 5% BSA in 1X lysis buffer for 1 hour, then blocking buffer discarded and beads washed thrice using 1X IP buffer (20 mM Tris-HCl pH 7.5, 150 mM KCl, 5 mM MgCl2 and 1 mM DTT), followed by incubation with specific antibody solution in 1X lysis buffer (final dilution 1:100) for 3 hours at 4⁰C. The cleared cell lysates, were incubated with primary antibody bound Protein-G Agarose bead (Invitrogen) and rotated overnight at 4⁰C. Thereafter, the beads were washed and then divided into two equal parts and each part was analyzed for bound proteins and RNAs by Western Blot and qRT-PCR respectively.

### Nanoparticle Tracking Analysis (NTA)

For characterization of EVs by NTA, the isolated EV pellet was resuspended in 1X PBS and 10-fold dilution was prepared to make 1ml suspension. This suspension was injected into the sample chamber of a nanoparticle tracker (Nanosight NS300, Malvern Panalytical). For characterization of isolated endosomes after *in vitro* assay, endosomal pellets were resuspended in 1ml 1X PBS and endosomal suspension was injected into the sample chamber of a nanoparticle tracker (Nanosight NS300, Malvern Panalytical).

### Atomic Force Microscopy

Endosomes were imaged and characterized by atomic force microscopy (AFM), and endosomal pellets were resuspended in autoclaved ultrapure water. Then, 5 µL of the endosomal suspension was spotted on a mica sheet (Muscovite Mica-V1, Electron Microscopy Sciences, Hatfield, PA) and dried for 15 minutes. If required, sample spots were gently washed with autoclaved ultrapure water to remove molecules that could interfere with imaging and then dried again. AAC mode of AFM was followed using a Pico plus 5500 ILM AFM (Agilent Technologies, USA) with a piezoscanner having maximum range of 9µm. Microfabricated silicon cantilevers having length of 225µm were used (Nano sensors, USA). Image processing was done using Picoview 1.1 software (Agilent Technologies, USA).

### Small RNA pool isolation

Small RNA pool from cellular lysate was isolated using miRVANA miRNA isolation kit (Thermofisher Scientific, USA) according to the manufacturer’s instructions. This small RNA pool was used as cargo for *in vitro* assay reactions and level of individual miRNA packaging was analyzed.

### In vitro mRNA transcription

*In vitro* transcription was performed by using mMESSAGE mMACHINE kit (Thermofisher Scientific, USA) and subsequently poly A tailing of transcripts were done with Poly (A) tailing kit (Thermofisher Scientific, USA) as per the manufacturer’s protocol. For preparation of linearized plasmid DNA template, RL-Con and RL-3X bulge-miR-122 were digested with DraI (New England Biolabs) prior to following the manufacturer’s protocol. The transcripts obtained were size verified by 6% Urea-PAGE and ethidium bromide-based detection and analysis.

### Phosphate 5’ end-labeling of synthetic miRNA

50 pmoles of synthetic miR-122 RNA (22nt) was incubated with T_4_ Polynucleotide Kinase (PNK) enzyme (10 units/µl) and 1mM ATP in 1X T4 PNK buffer at 37°C for 30 minutes at static condition. Reaction was ended with Tris-EDTA solution and was filtered through mini Quick spin oligo column (Roche Diagnostics). RNA was extracted with TriZol LS and CHCl_3_ and precipitated with isopropanol at -20°C overnight in presence of glycoblue co-precipitant. RNA pellet was resupended in nuclease-free water to a final concentration of 1 pmoles/ µl.

### Preparation of recombinant HuR

Recombinant HuR protein was purified by procedure as previously described in detail (Mukherjee et al., 2016). For expression of HuR in E. coli, the construct consisting HuR coding region in pET42a(+) was used. The protein was expressed in BL21DE3 *E. coli* cells codon Plus expression competent bacterial cells. IPTG induced overnight cultures of E. coli BL21 were lysed by incubation with lysis buffer [20 mM Tris–HCl, pH 7.5, 300 mM KCl, 2 mM MgCl2, 5 mM β-mercaptoethanol, 50 mM imidazole, 0.5% Triton X-100, 5% glycerol, 0.5 mg/ml lysozyme, 1× EDTA-free protease inhibitor cocktail (Roche)] for 30 min followed by sonication (10 seconds, three pulses). The cleared lysate was incubated with pre-equilibrated Ni-NTA Agarose beads (Qiagen) for 4 h at 4°C. Bead washing was done using wash buffer (20 mM Tris–HCl, pH 7.5,150 mM KCl, 2 mM MgCl2, 5 mM β-mercaptoethanol, 50 mM imidazole, 0.5% Triton X-100, 5% glycerol) with rotation, on an end-to-end rotator, each for 10 min, at 4°C. His-tagged HuR protein was eluted by incubating beads with different elution buffers (20 mM Tris–HCl, pH 7.5,150 mM KCl, 2 mM MgCl2, 5 mM β-mercaptoethanol, 0.5% Triton X-100, 5% glycerol) of different and increasing imidazole concentrations, each for 15 min at 4°C with end-to-end rotation. Purified protein was kept in aliquots and stored at −80°C.

### Free phosphate estimation assay for GTPase activity analysis

For estimation of RalA GTPase activity, free phosphate release in solution was analyzed by malachite green based colorimetric assay using a Phosphate assay kit (Sigma-Aldrich) by following manufacturer’s protocol. GTPase activity assay was performed with immobilized FH-RalA by addition of recombinant HuR and/or synthetic miRNA in presence of 1mM GTP at room temperature for 30 mins while shaking. After reaction, bead bound RalA was separated by centrifugation and released phosphate was present in supernatant due to cleavage of GTP by RalA GTPase activity. Tubes containing this supernatant were placed in ice and 50 µl from each set of samples were added in a 96 well clear plate on which 100 µl of malachite green reagent was added. The mix was incubated in dark for about 30 minutes for color to develop and absorbance at 620nm was measured and analyzed to study change in GTPase activity.

### Immunofluorescence and Confocal imaging

Cells were grown in 12 well plate on 18mm round gelatin coated coverslips and transfected as specified in Results. Cells were fixed using 4% paraformaldehyde (PFA) for 20 minutes at room temperature in dark condition. For indirect immunofluorescence, PFA fixed cells were blocked and permeabilized with PBS containing 1% BSA and 0.1% Triton X-100 and 10% goat serum (Gibco) for 30 minutes at room temperature. Coverslips were washed with 1X PBS thrice and then probed with fluorochrome tagged secondary antibodies (Alexa Fluor-conjugated, dilution 1:500) for 1 hour at room temperature. Zeiss LSM800 confocal microscope was used for imaging of fixed-cells. LSM800 was equipped with a Plan-Apo 63X/1.4NA oil immersion objective (Zeiss) and images were analyzed using Imaris7 software. Number and size of individual vesicles or bodies were measured by using particle generator of the Surpass plug-in available in Imaris7 software.

### Statistical Analysis

Software GraphPad Prism 5.00 (Graph Pad, San Diego) was used for analyzing the dataplots obtained from experiments performed at triplicate unless mentioned otherwise. Student’s t-test was done to determine P values. Significance was considered if P<0.05. Error bars represent mean±s.d.

### Data Availability

All data supporting the findings of this study are available from the corresponding authors upon request.

### Conflict of Interest

The authors declare no conflict of interest

## Acknowledgements

We thank Witold Filipowicz and Gunter Meister for different constructs used in this study. We thank the Funding body, Dept. of Science and Technology (DST), Govt. of India along with Council for Scientific and Industrial Research (CSIR), and University Grant Commission (UGC) for the fellowship to SG and SHC respectively. SNB is supported by The Swarnajayanti Fellowship (DST/SJF/LSA-03/2014-15) from Dept. of Science and Technology, Govt. of India. The work also received support from a High-Risk High Reward Grant (HRR/2016/000093) from Dept. of Science and Technology, Govt. of India and CEFIPRA project grant 6003-J. SNB Also acknowledge the Start-Up Support Grant of University of Nebraska, USA.

## Author Contributions

S.N.B. and K.M. conceived the idea, designed the experiments and analysed the data. S.G. S.H.C have contributed in design and planning the experiments. S.G. S.H.C performed the experiments. K.M. and S.G. also wrote the manuscript with S.N.B. and analysed the data.

## Supplementary Information

**Figure S1:**
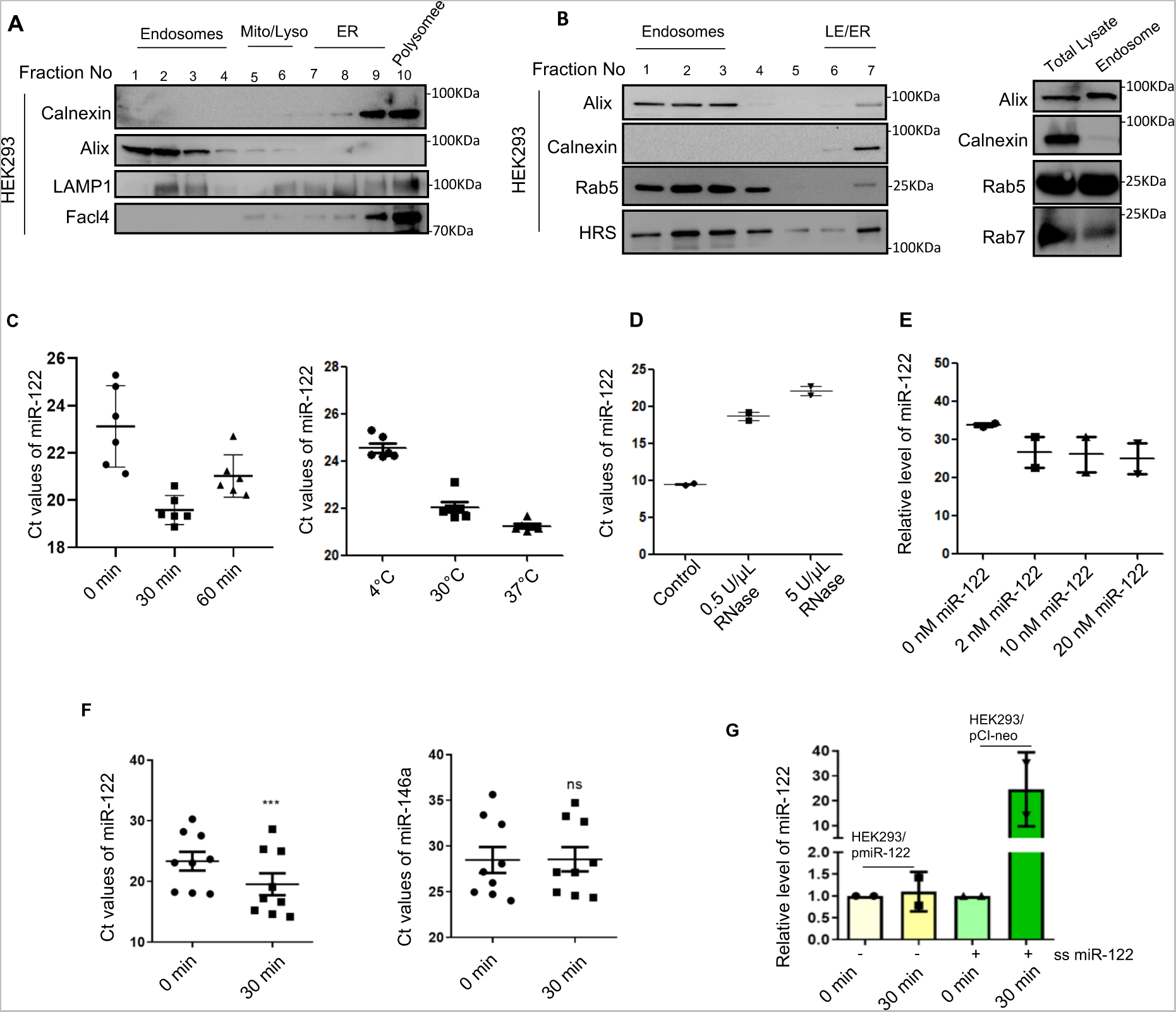
Initial Cell-free reactions using HEK293 endosomes and Normalization of miRNA packaging in C6 endosomes. **A.** Immunoblots of various cellular marker proteins in different fractions obtained from 3-30% iodixanol gradient after subcellular fractionation of HEK293 post-nuclear lysate by ultracentrifugation. **B.** Western blots of marker proteins in different fractions obtained from 3-15% iodixanol step gradient after fractionation of HEK293 post-nuclear lysate for endosome enrichment and also the immunoblots comparing total cellular lysate with enriched endosomal lysate. Endosomal pellet fraction is free from ER contamination as observed from calnexin blots. Enriched for early endosomes, indicated by presence of Rab5 and Rab7. **C.** Import of miR-122 into endosomes in the in vitro assay done with HEK293 endosomes using synthetic miR-122 as cargo under different incubation time and temperature of the reaction. After the reaction, endosomes were trated with RNase and protected miRNA was recovered, and RT-PCR analysis was performed. Ct values of the miR-122 were plotted. **D.** RNase A protection assay and effective concentration of RNase A for degradation of miRNA not packaged into endosomes. Two different concentrations of RNA were used to digest the non-incorporated miRNAs before the endosomes were recovered and miR-122 content was measured after the reaction of 30 minutes at 37°C. **E.** Concentration dependnent increase of mR-122 import in vitro with HEK293 endosomes. After the incubation with increasing concentration of substrate miRNA, The RNase protected miRNA was recovered, and RT-PCR analysis was performed. Ct values of the miR-122 were plotted. **F.** Change in Ct values in imported miR-122 and endogenously present miR-146a after the HEK293 endosmes were incubated with miR-122 as substrate RNA for 0 and 30 mins. RNA was recovered, and RT-PCR analysis was performed (n= 3 independent experiments for both panels, P= 0.0001 and 0.7128 for left and right panels respectively). **G.** Cell-free assay with endosomes isolated from miR-122 expressing C6 cells in presence and absence of additional synthetic miR-122 as substrate for 0 and 30 min in vitro before the RNA was reisolated after RNase treatment. Recovered RNA was analyzed for miR-122 content. miR-146a was used as internal control and relative value of miR-122 was normalized to that. Data information: In all the experimental data, error bars are represented as mean with SD, ns, nonsignificant, ***P < 0.001, respectively. P-values were calculated by two-tailed paired t-test in most of the experiments unless mentioned otherwise. Relative level of miR-122 was normalized with miR-146a level by 2^-ΔΔCt^ method. Positions of molecular weight markers are marked and shown with the respective Western blots.

**Figure S2:**
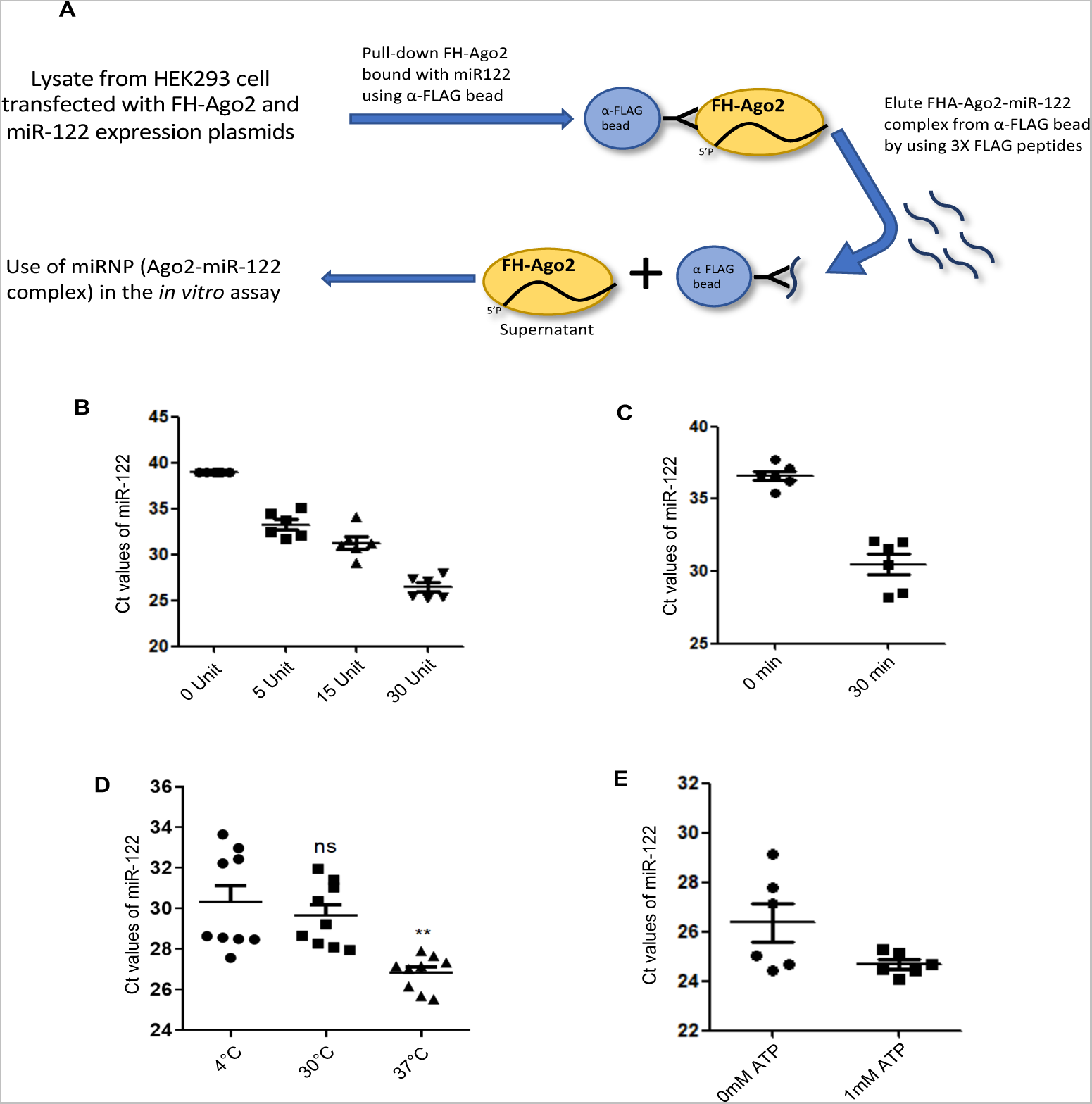
Isolated endosomes able to package miRNA from Ago2 miRNP complex in cell-free reaction. **A.** Schematic work-flow of isolation of miRNA bound with Ago2 protein (miRNP) complex from HEK293 cells transfected with pre-miR-122 expression plasmid and FH-Ago2 to get Ago2 miRNP as cargo in cell-free reaction to study their packaging into endosomes. **B.** Optimization of miRNP concentration as cargo in the assay. After the reaction, endosomes were trated with RNase and protected miRNA was recovered, and RT-PCR analysis was performed. Ct values of the miR-122 were plotted. **C.** Time dependent increase in RNase protected miRNA in endosomes after 30 min of incubation with 30 units of miR-122 miRNPs *in vitro.* After the reaction, endosomes were trated with RNase and protected miRNA was recovered, and RT-PCR analysis was performed. Ct values of the miR-122 were plotted. **D.** Effect of temperature on miRNA imported when miR-122 miRNPs was used as substrate. Reaction was carried out at different temperature and after the reaction, endosomes were trated with RNase and protected miRNA was recovered, and RT-PCR analysis was performed. Ct values of the miR-122 were plotted. (n=3 independent experiments, P= 0.3763, 0.0078). **E.** miRNA packaging into endosome is facilitated by presence of ATP. miR-122 miRNPs was used as substrate. Reaction was carried out in presence and absence of ATP and after the reaction, endosomes were trated with RNase and protected miRNA was recovered, and RT-PCR analysis was performed. Ct values of the miR-122 were plotted. Data information: In all the experimental data, error bars are represented as mean with SD, ns, nonsignificant, **P < 0.01, respectively. P-values were calculated by two-tailed paired t-test.

**Figure S3:**
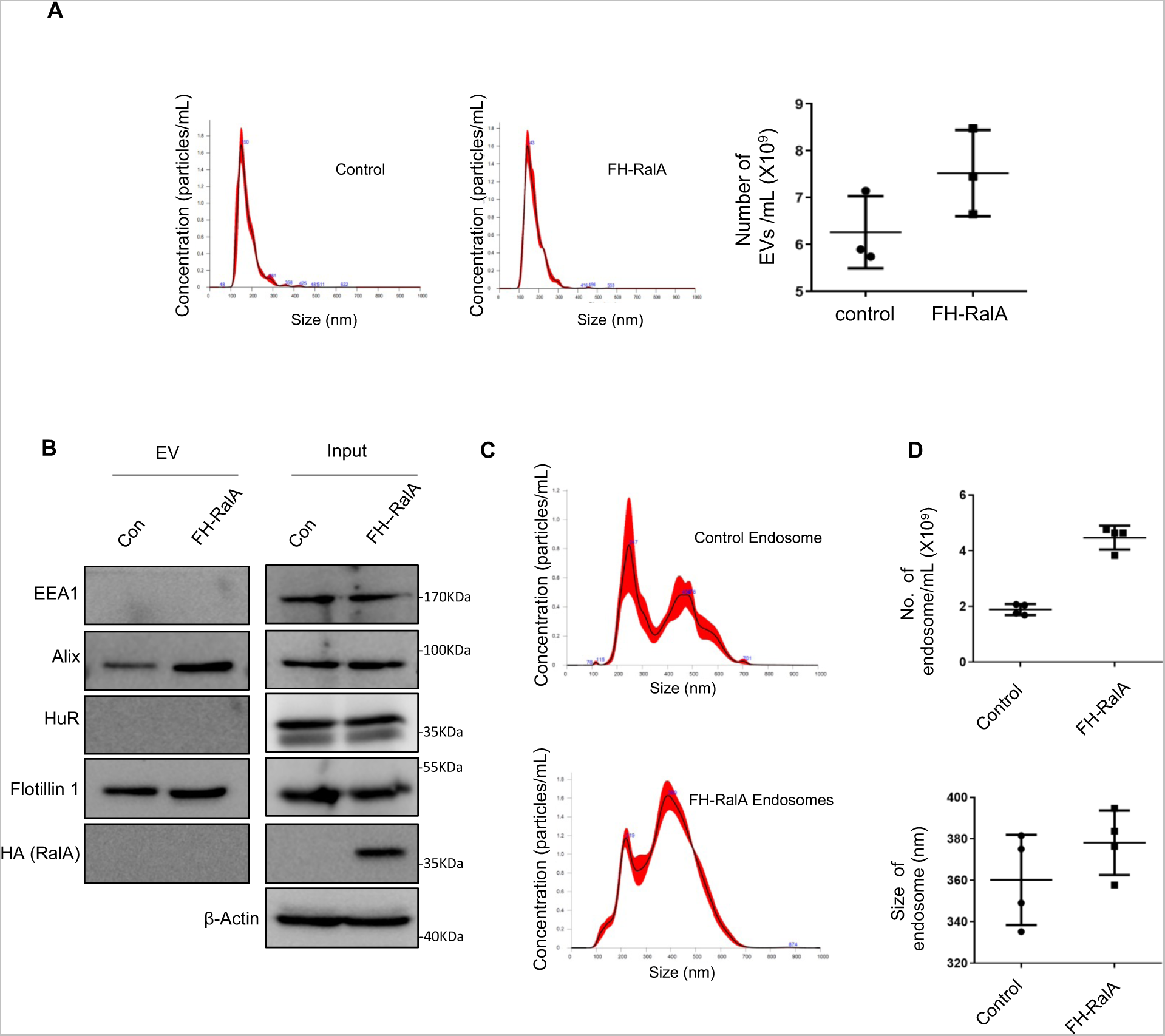
Effect of HA-RalA expression on extracellular vesicles (EVs) and endosomes. **A.** Nanoparticle tracking analysis (NTA) of EVs isolated from cells expressing pCI-Neo (control) or FH-RalA plasmids. FH-RalA expression lead to increase in secretion of EVs as can be seen from the number of EVs plotted in the graph. **B.** Western blots of different markers proteins done with EVs isolated and respective cellular samples. **C.** NTA analysis histograms of endosomes isolated from control and FH-RalA expressing cells. **D.** Estimation of size and number of endosomes showed increase in both due to expression of FH-RalA. Data are from three replicates.

**Figure S4.**
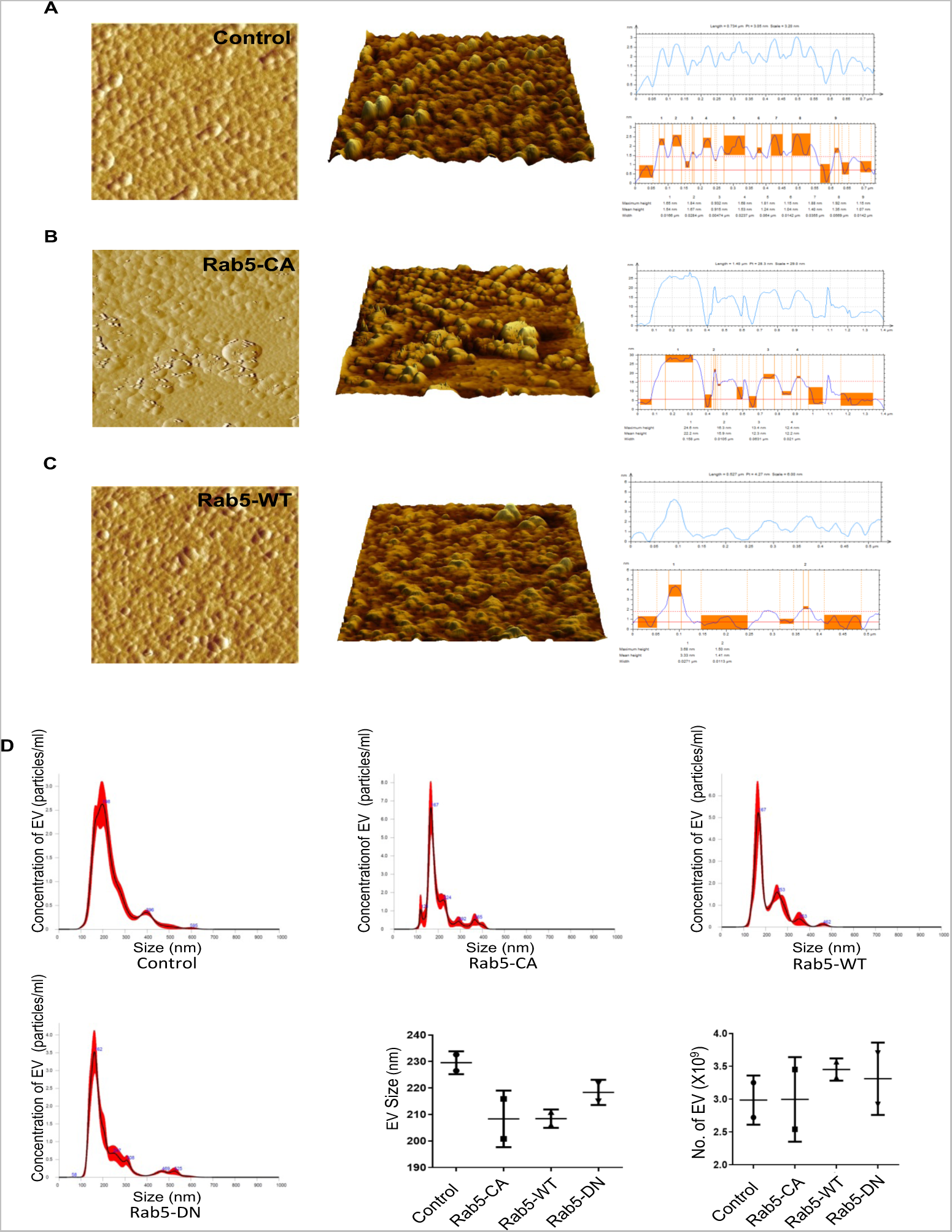
Structural analysis of endosomes derived from cells expressing wild type Rab5 and Rab5-CA mutant by Atomic Force Microscopy. Atomic Force Microscopy (AFM) of endosomes isolated from C6 cells expressing pCI-Neo (control), Rab5-CA and Rab5-WT plasmids. AFM micrographs depict 2D, 3D and amplitude plots (left to right) of **A.** Control cell derived endosomes **B.** Rab5-CA expressing cell derived endosomes **C.** Rab5-WT expressing cell derived endosomes. **D.** Nanoparticle tracking analysis (NTA) of EVs isolated from C6 cells expressing pCI-Neo (control), Rab5-CA, Rab5-DN and Rab5-WT plasmids.Size and numver of EVs are depicted in associated graphs. Data are from three replicates.

**Figure S5.**
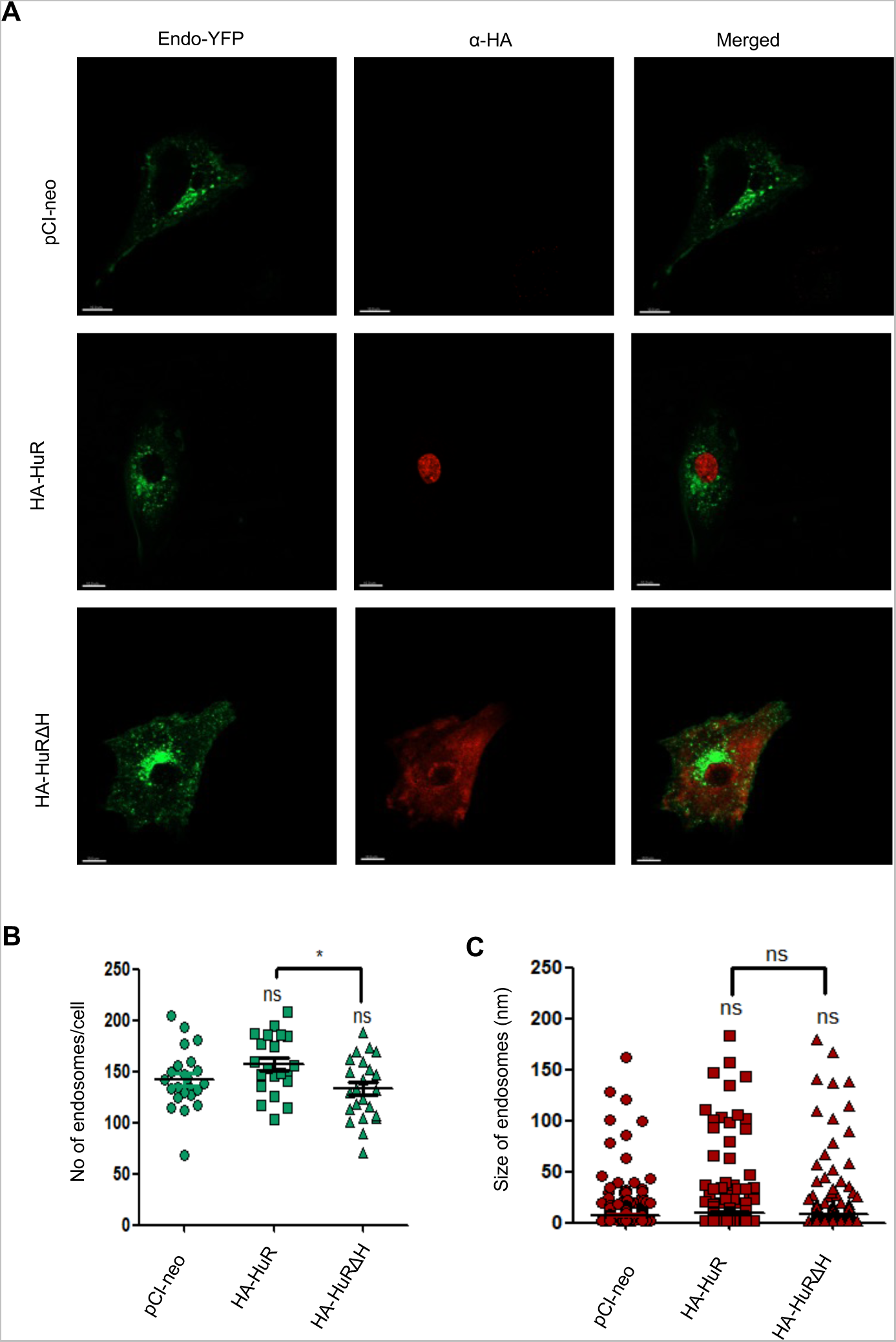
Effect of expression of HuR and Rab5 proteins on endosomal structures. **A.** Confocal images of C6 cells expressing of HA-HuR and the mutant HA-HuRΔH along with Endo-YFP. Endosomes were tagged by Endo-YFP (green) and expression of HA-HuR and HA-HuRΔH were indirectly visualized by using α-HA based immunofluorescence. **B-C**. Number and size of endosomes were analyzed by using IMARIS software (B panel of endosome number- n≥21 number of fields, P= 0.0865, 0.166, unpaired t-test P= 0.0103; C panel of endosome size- n≥80 number of endosomes, P= 0.0907, 0.3571, unpaired t-test P= 0.6441). Scale bar 10 µM. Data information: In all the experimental data, error bars are represented as mean with SD, ns, nonsignificant, *P < 0.05, respectively. P-values were calculated by two-tailed paired t-test in most of the experiments unless mentioned otherwise.

**Supplementary Table 1:** List of Plasmids.

| **Name of plasmids** | **Reference/Source** | **Plasmid description** |
| --- | --- | --- |
| pmiR-122 | Received from Chang J. | Plasmid encoding pre-miR-122 under a constitutive U6 promoter |
| pmiR-146a | Kind gift of Dr. Susanta Roychoudhury | miR-146a expressing pU61 construct |
| HA-HuR and HA-HuRΔH | Kind gift of Witold Filipowicz | Full length and truncated HuR without hinge region (amino acids 186-242 deleted) with HA coding sequence in pCI-Neo vector |
| Flag-HA-Ago2 | Kind gift of Tom Tuschl | Expressing FLAG & HA tagged human Ago2 |
| Flag-HA-RalA | Constructed by cloning | RalA coding sequence was sub-cloned into Flag-HA-Ago2 vector between the NotI and BamHI sites replacing Ago2. Flag and HA tag at the N-terminal of RalA. |
| Flag-HA-Stx5 | Constructed by cloning | Stx5 coding sequence was sub-cloned into Flag-HA-Ago2 vector between the NotI and BamHI sites replacing Ago2. Flag and HA tag at the N-terminal of Stx5. |
| Myc-HuR | From Witold Filipowicz | HuR coding sequence between the EcoRI and XhoI sites of pcDNA3.1 containing the myc tag sequence between BamHI and EcoRI sites |
| Endo-YFP | Kind gift from Edouard Bertrand | Yellow Fluorescent Protein (YFP) tagged early endosome expression plasmid |
| Rab5-CA | Kind gift from J. M. Backer. | A GTPase-deficient mutant of Rab5a, Rab5Q79L expression plasmid |
| Rab5-WT | Kind gift from J. M. Backer. | Wild type human Rab5a expression plasmid |
| Rab5-DN | Kind gift from J. M. Backer. | A GTP-binding defective mutant of Rab5a, Rab5S34N expression plasmid |

**Supplementary Table 2:**
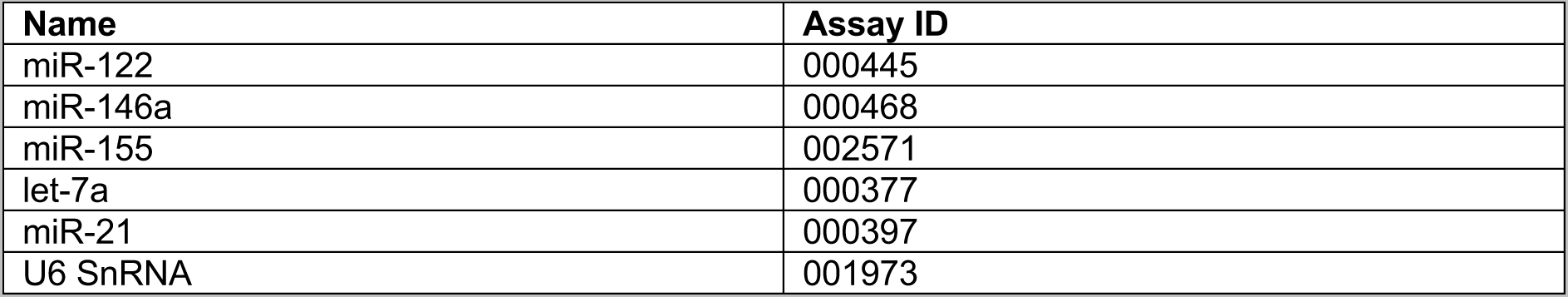
Details of miRNA primers used for Taqman based quantification.

| **Name** | **Assay ID** |
| --- | --- |
| miR-122 | 000445 |
| miR-146a | 000468 |
| miR-155 | 002571 |
| let-7a | 000377 |
| miR-21 | 000397 |
| U6 SnRNA | 001973 |

**Supplementary Table 3:** Details of Antibodies used for western blot (WB), Immunofluorescence (IF) and immunoprecipitation (IP)

| Antibody Name | Raised in | Dilution | Source | Catalogue No. |
| --- | --- | --- | --- | --- |
| Alix | Mouse | WB-1:100 | Santa Cruz | sc53538 |
| Calnexin | Rabbit | WB-1:10,000 | Bethyl | A303-696A |
| HRS | Rabbit | WB-1:2000 | Bethyl | A300-989A |
| Rab5a | Rabbit | WB-1:1000<br>IF- 1:100 | Cell Signaling | #3547S |
| Rab7a | Rabbit | WB-1:1000 | Cell Signaling | #9367S |
| EEA1 | Rabbit | WB-1:1000 | Cell Signaling | #2411S |
| Lamp1 | Rabbit | WB-1:1000 | Cell Signaling | #3243S |
| FacI4 | Rabbit | WB-1:1000 | Novus Biologicals | NBP2-16401 |
| HuR | Mouse | WB-1:1000 | Santa Cruz | sc-5261 (3A2) |
| HA | Rat | WB-1:1000<br>IP- 1:100<br>IF:100 | Roche | 11867431001 |
| β-Actin | Mouse monoclonal (HRP-conjugated) | WB-1:10,000 | Sigma Aldrich | A3854 |
| c-Myc | Mouse | WB-1:500 | Santa Cruz | Sc-40 (9E10) |
| RalA | Rabbit | WB-1:1000 | Cell Signaling | #3526 |
| Stx5 | Rabbit | WB-1:10,000 | Bethyl | A305-382A |
| Flotilin-1 | Rabbit | WB-1:1000 | Cell Signaling | #18634S |
| Ago2 (eIF2C2) | Mouse | WB-1:500 | Abnova | H00027161-M01 |

**Supplementary Table 4:**
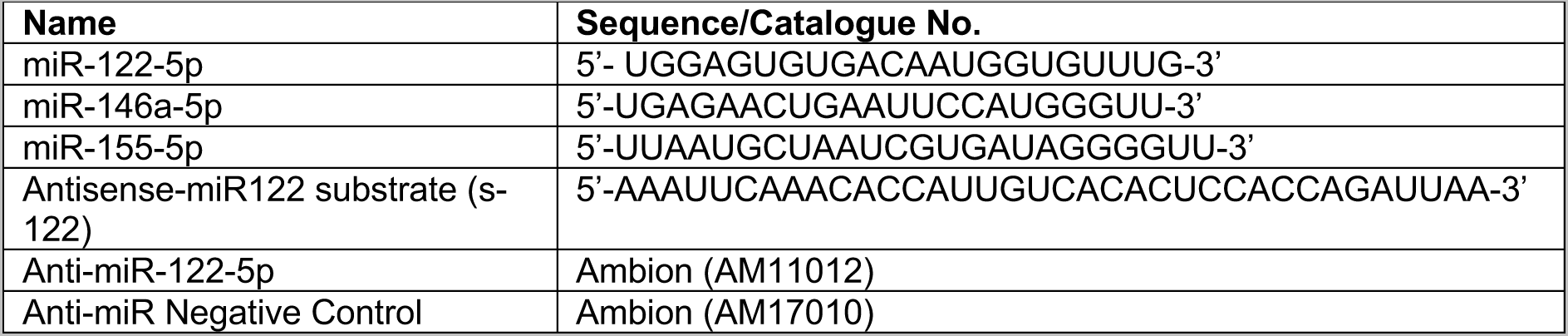
List of Synthetic single stranded RNA and anti-miR oligonucleotides.

## Notes

### Competing Interest Statement

The authors have declared no competing interest.

## References

Arroyo, J.D., J.R. Chevillet, E.M. Kroh, I.K. Ruf, C.C. Pritchard, D.F. Gibson, P.S. Mitchell, C.F. Bennett, E.L. Pogosova-Agadjanyan, D.L. Stirewalt, J.F. Tait, and M. Tewari. 2011. Argonaute2 complexes carry a population of circulating microRNAs independent of vesicles in human plasma. Proc Natl Acad Sci U S A. 108:5003–5008.

Bartel, D.P. 2009. MicroRNAs: target recognition and regulatory functions. Cell. 136:215–233.

Bhattacharyya, S.N., and S. Adhya. 2004. tRNA-triggered ATP hydrolysis and generation of membrane potential by the leishmania mitochondrial tRNA import complex. The Journal of biological chemistry. 279:11259–11263.

Bose, M., S. Chatterjee, Y. Chakrabarty, B. Barman, and S.N. Bhattacharyya. 2020. Retrograde trafficking of Argonaute 2 acts as a rate-limiting step for de novo miRNP formation on endoplasmic reticulum-attached polysomes in mammalian cells. Life science alliance. 3.

Bucci, C., R.G. Parton, I.H. Mather, H. Stunnenberg, K. Simons, B. Hoflack, and M. Zerial. 1992. The small GTPase rab5 functions as a regulatory factor in the early endocytic pathway. Cell. 70:715–728.

Buschow, S.I., B.W. van Balkom, M. Aalberts, A.J. Heck, M. Wauben, and W. Stoorvogel. 2010. MHC class II-associated proteins in B-cell exosomes and potential functional implications for exosome biogenesis. Immunol Cell Biol. 88:851–856.

De, D., and S.N. Bhattacharyya. 2021. Amyloid-beta oligomers block lysosomal targeting of miRNPs to prevent miRNP recycling and target repression in glial cells. Journal of cell science. 134.

De, D., I. Mukherjee, S. Guha, R.K. Paidi, S. Chakrabarti, S.C. Biswas, and S.N. Bhattacharyya. 2021. Rheb-mTOR activation rescues Abeta-induced cognitive impairment and memory function by restoring miR-146 activity in glial cells. Mol Ther Nucleic Acids. 24:868–887.

Flinn, R.J., Y. Yan, S. Goswami, P.J. Parker, and J.M. Backer. 2010. The late endosome is essential for mTORC1 signaling. Molecular biology of the cell. 21:833–841.

Garcia-Martin, R., G. Wang, B.B. Brandao, T.M. Zanotto, S. Shah, S. Kumar Patel, B. Schilling, and C.R. Kahn. 2022. MicroRNA sequence codes for small extracellular vesicle release and cellular retention. Nature. 601:446–451.

Ghosh, J., M. Bose, S. Roy, and S.N. Bhattacharyya. 2013. Leishmania donovani targets Dicer1 to downregulate miR-122, lower serum cholesterol, and facilitate murine liver infection. Cell Host Microbe. 13:277–288.

Ghosh, S., M. Bose, A. Ray, and S.N. Bhattacharyya. 2015. Polysome arrest restricts miRNA turnover by preventing exosomal export of miRNA in growth-retarded mammalian cells. Molecular biology of the cell. 26:1072–1083.

Ghosh, S., K. Mukherjee, Y. Chakrabarty, S. Chatterjee, B. Ghoshal, and S.N. Bhattacharyya. 2021. GW182 Proteins Restrict Extracellular Vesicle-Mediated Export of MicroRNAs in Mammalian Cancer Cells. Molecular and cellular biology. 41.

Ghoshal, B., E. Bertrand, and S.N. Bhattacharyya. 2021. Non-canonical ago loading of EV-derived exogenous single stranded miRNA in recipient cells. Journal of cell science.

Gorvel, J.P., P. Chavrier, M. Zerial, and J. Gruenberg. 1991. rab5 controls early endosome fusion in vitro. Cell. 64:915–925.

Goswami, A., K. Mukherjee, A. Mazumder, S. Ganguly, I. Mukherjee, S. Chakrabarti, S. Roy, S. Sundar, K. Chattopadhyay, and S.N. Bhattacharyya. 2020. MicroRNA exporter HuR clears the internalized pathogens by promoting pro-inflammatory response in infected macrophages. EMBO Mol Med. 12:e11011.

Groot, M., and H. Lee. 2020. Sorting Mechanisms for MicroRNAs into Extracellular Vesicles and Their Associated Diseases. Cells. 9.

Hammond, S.M. 2015. An overview of microRNAs. Adv Drug Deliv Rev. 87:3–14.

Hurley, J.H. 2010. The ESCRT complexes. Crit Rev Biochem Mol Biol. 45:463–487.

Hyenne, V., A. Apaydin, D. Rodriguez, C. Spiegelhalter, S. Hoff-Yoessle, M. Diem, S. Tak, O. Lefebvre, Y. Schwab, J.G. Goetz, and M. Labouesse. 2015. RAL-1 controls multivesicular body biogenesis and exosome secretion. The Journal of cell biology. 211:27–37.

Iavello, A., V.S. Frech, C. Gai, M.C. Deregibus, P.J. Quesenberry, and G. Camussi. 2016. Role of Alix in miRNA packaging during extracellular vesicle biogenesis. Int J Mol Med. 37:958–966.

Janas, M.M., B. Wang, A.S. Harris, M. Aguiar, J.M. Shaffer, Y.V. Subrahmanyam, M.A. Behlke, K.W. Wucherpfennig, S.P. Gygi, E. Gagnon, and C.D. Novina. 2012. Alternative RISC assembly: binding and repression of microRNA-mRNA duplexes by human Ago proteins. RNA. 18:2041–2055.

Kahya, N., E.I. Pecheur, W.P. de Boeij, D.A. Wiersma, and D. Hoekstra. 2001. Reconstitution of membrane proteins into giant unilamellar vesicles via peptide-induced fusion. Biophys J. 81:1464–1474.

Kosaka, N., H. Iguchi, Y. Yoshioka, F. Takeshita, Y. Matsuki, and T. Ochiya. 2010. Secretory mechanisms and intercellular transfer of microRNAs in living cells. J Biol Chem. 285:17442–17452.

Lang, C.H., H.L. Blakesley, G.J. Bagby, and J.J. Spitzer. 1988. Lipoxygenase and cyclooxygenase blockade by BW755C prevents endotoxin-induced hypotension but not changes in glucose metabolism. Horm Metab Res. 20:551–554.

Lebrand, C., M. Corti, H. Goodson, P. Cosson, V. Cavalli, N. Mayran, J. Faure, and J. Gruenberg. 2002. Late endosome motility depends on lipids via the small GTPase Rab7. The EMBO journal. 21:1289–1300.

Lee, S.H., and C. Mayr. 2019. Gain of Additional BIRC3 Protein Functions through 3’-UTR-Mediated Protein Complex Formation. Mol Cell. 74:701–712 e709.

Linders, P.T., C.V. Horst, M.T. Beest, and G. van den Bogaart. 2019. Stx5-Mediated ER-Golgi Transport in Mammals and Yeast. Cells. 8.

Liu, A.P., and D.A. Fletcher. 2009. Biology under construction: in vitro reconstitution of cellular function. Nat Rev Mol Cell Biol. 10:644–650.

Liu, X.M., L. Ma, and R. Schekman. 2021. Selective sorting of microRNAs into exosomes by phase-separated YBX1 condensates. Elife. 10.

Mazan-Mamczarz, K., P.R. Hagner, S. Corl, S. Srikantan, W.H. Wood, K.G. Becker, M. Gorospe, J.D. Keene, A.S. Levenson, and R.B. Gartenhaus. 2008. Post-transcriptional gene regulation by HuR promotes a more tumorigenic phenotype. Oncogene. 27:6151–6163.

McKenzie, A.J., D. Hoshino, N.H. Hong, D.J. Cha, J.L. Franklin, R.J. Coffey, J.G. Patton, and A.M. Weaver. 2016. KRAS-MEK Signaling Controls Ago2 Sorting into Exosomes. Cell Rep. 15:978–987.

Mukherjee, K., B. Ghoshal, S. Ghosh, Y. Chakrabarty, S. Shwetha, S. Das, and S.N. Bhattacharyya. 2016. Reversible HuR-microRNA binding controls extracellular export of miR-122 and augments stress response. EMBO reports. 17:1184–1203.

Muralidharan-Chari, V., J.W. Clancy, A. Sedgwick, and C. D’Souza-Schorey. 2010. Microvesicles: mediators of extracellular communication during cancer progression. Journal of cell science. 123:1603–1611.

Nabhan, J.F., R. Hu, R.S. Oh, S.N. Cohen, and Q. Lu. 2012. Formation and release of arrestin domain-containing protein 1-mediated microvesicles (ARMMs) at plasma membrane by recruitment of TSG101 protein. Proc Natl Acad Sci U S A. 109:4146–4151.

Raiborg, C., and H. Stenmark. 2009. The ESCRT machinery in endosomal sorting of ubiquitylated membrane proteins. Nature. 458:445–452.

Raposo, G., and W. Stoorvogel. 2013. Extracellular vesicles: exosomes, microvesicles, and friends. J Cell Biol. 200:373–383.

Santangelo, L., G. Giurato, C. Cicchini, C. Montaldo, C. Mancone, R. Tarallo, C. Battistelli, T. Alonzi, A. Weisz, and M. Tripodi. 2016. The RNA-Binding Protein SYNCRIP Is a Component of the Hepatocyte Exosomal Machinery Controlling MicroRNA Sorting. Cell Rep. 17:799–808.

Shurtleff, M.J., M.M. Temoche-Diaz, K.V. Karfilis, S. Ri, and R. Schekman. 2016. Y-box protein 1 is required to sort microRNAs into exosomes in cells and in a cell-free reaction. Elife. 5.

Squadrito, M.L., C. Baer, F. Burdet, C. Maderna, G.D. Gilfillan, R. Lyle, M. Ibberson, and M. De Palma. 2014. Endogenous RNAs modulate microRNA sorting to exosomes and transfer to acceptor cells. Cell Rep. 8:1432–1446.

Stuffers, S., C. Sem Wegner, H. Stenmark, and A. Brech. 2009. Multivesicular endosome biogenesis in the absence of ESCRTs. Traffic. 10:925–937.

Thery, C., S. Amigorena, G. Raposo, and A. Clayton. 2006. Isolation and characterization of exosomes from cell culture supernatants and biological fluids. Curr Protoc Cell Biol. Chapter 3:Unit 3 22.

Trajkovic, K., C. Hsu, S. Chiantia, L. Rajendran, D. Wenzel, F. Wieland, P. Schwille, B. Brugger, and M. Simons. 2008. Ceramide triggers budding of exosome vesicles into multivesicular endosomes. Science. 319:1244–1247.

Villarroya-Beltri, C., C. Gutierrez-Vazquez, F. Sanchez-Cabo, D. Perez-Hernandez, J. Vazquez, N. Martin-Cofreces, D.J. Martinez-Herrera, A. Pascual-Montano, M. Mittelbrunn, and F. Sanchez-Madrid. 2013. Sumoylated hnRNPA2B1 controls the sorting of miRNAs into exosomes through binding to specific motifs. Nat Commun. 4:2980.

Xie, Y., and Y. Ren. 2019. Mechanisms of nuclear mRNA export: A structural perspective. Traffic. 20:829–840.

